# Combining Machine Learning and Directed Evolution for Optimization of a Monooxygenase

**DOI:** 10.64898/2026.07.31.742090

**Authors:** Daniel Gutierrez, Isa Madrigal Harrison, Aaron Feller, Andrew D. Ellington

## Abstract

L-3,4-dihydroxyphenylalanine (L-Dopa) is an important pharmaceutical for the treatment of Parkinson’s disease and a precursor to numerous catechol-containing compounds. The flavin-dependent monooxygenase HpaBC is a promising biocatalyst for microbial L-Dopa production but exhibits limited native activity toward L-tyrosine. Although structure-based machine learning (ML) models have become increasingly popular for protein engineering, relatively few studies have systematically compared their performance or evaluated their integration into iterative engineering workflows. Here, we benchmarked multiple ML models for their ability to predict activity enhancing mutations in HpaBC. Experimentally validated single mutants were used to seed combinatorial design with EVOLVEpro, generating progressively improved higher-order variants. We next evaluated how expanding the EVOLVEpro training set with directed evolution derived variants influenced combinatorial predictions and finally explored an expanded sequence space by allowing combinations of both machine learning derived and directed evolution derived mutations. This workflow produced HpaBC variants with substantially improved activity. Although incorporating directed evolution data substantially altered EVOLVEpro’s predicted mutational trajectories, both training strategies converged on variants with comparable activities, demonstrating that distinct regions of sequence space can yield similarly optimized enzymes. Together, these results provide a systematic comparison of zero-shot ML models and establish an iterative framework for integrating machine learning with directed evolution to accelerate enzyme engineering.

## Introduction

L-3,4-dihydroxyphenylalanine (L-Dopa) is a high-value aromatic amino acid that is widely used as a first-line treatment for Parkinson’s disease^1,2^. Beyond its clinical importance, L-Dopa is a versatile precursor for the biosynthesis of numerous catechol-containing molecules, including neurotransmitters such as dopamine, norepinephrine, and epinephrine. These catechol-containing compounds have applications spanning medicine, cosmetics, functional materials, and natural product synthesis^3–5^. Hence, generating improved pathways for L-Dopa may prove especially useful both for creating coatings and for developing robust pathways for the high-titer production of biomedically useful alkaloids.

In this regard, the enzyme 4-hydroxyphenylacetate 3-monooxygenase (HpaBC) has proven to be an excellent workhorse for the production of L-Dopa^6^. HpaBC is a two-component enzyme system consisting of the FAD-dependent monooxygenase HpaB and the NAD(P)H-dependent flavin reductase HpaC^7^. HpaB catalyzes the hydroxylation reaction, while HpaC regenerates the reduced flavin cofactor by transferring electrons from NAD(P)H to FAD, supplying the FADH₂ required for HpaB activity^8^. Previous directed evolution campaigns have led to HpaBC variants that produce 45x more L-Dopa then the wild-type enzyme^9^.

We wanted to determine whether machine learning methods could be used to further improve production of L-Dopa by HpaBC. Recent advances in machine learning offer an attractive alternative to directed evolution for protein engineering^10–17^. By prioritizing beneficial substitutions prior to experimental validation, the size of the experimental search space is greatly reduced; random walks become less necessary or useful.

In particular, structure-based deep learning models have emerged as promising tools^18–20^, and existing structure-based methods span a range of approaches, from models that evaluate local residue environments^18^ to generative approaches that redesign entire protein sequences while preserving structural compatibility^20^. In parallel, protein language models, such as Evolutionary Scale Modeling (ESM)^21^, leverage the evolutionary information inherent in protein sequence databases and can be used to predict intrinsic biological properties as emergent behaviors^22^.

However, because of the large number of possible approaches, it is difficult for engineers to determine which may be most effective. In this study, we evaluated a number of machine learning models for their ability to predict activity enhancing substitutions in HpaBC. In general, each model was similarly successful, but predictions varied. We leveraged EVOLVEpro^23^ to evaluate multi-mutant variants that were found to be as active as those obtained by directed evolution. We find that the most active variants arose either from directed evolution or included variants from directed evolution as starting points for further engineering.

## Materials and Methods

### Alphafold2 ColabFold

Folded structures for zero-shot predictions were prepared using the ColabFold implementation of Alphafold 2^24–26^ (using the sequence from an *E. coli* HpaB pdb:6EB0^8^) in monomeric and functional tetrameric form for use in zero-shot prediction with MutCompute and ProteinMPNN. It should be noted that we used the 6EB0 from the PDB for structure^8^, which differs slightly (by two amino acids at the carboxy terminus; N478S,D499N) from the sequence of the protein that had previously been proofed in our assay format^9^.

### MutCompute Zero-shot Prediction

MutCompute^18^ was run using the above monomeric and tetrameric structures. Scores for each variant were calculated by dividing the mutant probability for each residue by the wildtype residue probability. Variants were then ranked in descending order, and the top 60 mutants were selected for both monomeric and tetrameric structures.

### ProteinMPNN Zero-shot Prediction

ProteinMPNN^20^ was run using the above monomeric and tetrameric structures at a temperature of .1 saving the design statistics. Scores for each variant were calculated by dividing the mutant probability for each residue by the wildtype residue probability from the design probability matrix. Variants were then ranked in descending order, and the top 60 mutants were selected for both monomeric and tetrameric structures.

### ESM Zero-shot Prediction

ESM Zero Shot Prediction was performed as in Meier et. al.^27^ using the masked-marginals setting and sequence from HpaB 6EB0. This was done for ESM1V, ESM2 650M, and ESM2 15B^28^, the top 60 mutants were selected via this scoring metric for each version of ESM.

### Bacterial strains and growth conditions

Unless otherwise stated, all experiments were performed in *Escherichia coli* DH10B (New England Biolabs). Cultures were grown in LB medium (Lennox formulation; Fisher BioReagents) supplemented with 100 μg mL⁻¹ carbenicillin and/or 34 μg mL⁻¹ chloramphenicol, as appropriate for plasmid selection.

### Competent cell preparation

An *E. coli* DH10B reporter strain carrying the DopA reporter plasmid was generated by chemical transformation. The resulting strain was made chemically competent using the Mix & Go *E. coli* Transformation Buffer Set (Zymo Research) according to the manufacturer’s protocol, using SOB medium prepared in-house as described by Zymo Research. These competent reporter cells were used for all subsequent cloning, transformations, and variant screening experiments.

### Plasmid construction

Plasmids were assembled by Golden Gate cloning using BsaI-HFv2 (New England Biolabs). Plasmid sequences and complete construct sequences are provided in Supplementary Table 1 and 2. Synthetic DNA fragments were obtained from Twist Bioscience.

### Chemical transformation

Golden Gate assembly reactions were transformed into chemically competent DH10B reporter cells according to the Mix & Go (Zymo Research) protocol. Following recovery in SOC medium, transformants were plated on LB agar (Miller formulation) supplemented with the appropriate antibiotics and incubated overnight at 37°C.

### L-Dopa biosensor assay

Three independent colonies from each construct were inoculated into 1 mL of LB medium containing the appropriate antibiotics in 96-well deep-well plates and grown overnight (approximately 16 h) at 37°C with shaking at 950 rpm.

The following day, overnight cultures were diluted (1:50) into fresh LB medium containing the appropriate antibiotics and 0.5 mg mL⁻¹ L-ascorbic acid and grown for 2 h at 37°C with shaking at 950 rpm. For each biological replicate, paired induced and uninduced cultures were prepared. Induced cultures were supplemented with anhydrotetracycline (aTc) to a final concentration of 20 nM, while paired cultures remained uninduced. Cultures were incubated for an additional 4 h under the same conditions.

Following induction, aliquots of each culture were transferred to black, clear-bottom 96-well microplates (Corning, Cat. No. 3631). Optical density at 600 nm (OD600) and GFP fluorescence (excitation, 485 nm; emission, 535 nm) were measured using a Tecan Infinite M Plex.

Fluorescence values were normalized to cell density (RFU/OD600). For each biological replicate, background fluorescence from the corresponding uninduced culture was subtracted from the induced fluorescence value. Normalized fluorescence values were subsequently expressed relative to the wild-type control assayed on the same plate.

### EVOLVEpro

EVOLVEpro was used once to rank all combinatorial double, triple, quadruple, and quintuple mutants consisting only of single mutants exhibiting a 1.25X fold change in fluorescence as measured in our assay. The top 5 double, triple, quadruple, and quintuple mutants were selected, cloned, transformed, then assayed in the L-Dopa biosensor assay described above. A second round of EVOLVEpro variants were selected from the same set of mutations but included all combinatorial hextuple mutants of which the top 6 were selected and assayed.

EVOLVEpro was trained with additional combinatorial mutant data from Gilmore et. al.^9^ and ranking the same set of hextuple mutants of which the top 6 were selected and assayed.

EVOLVEpro was provided variants that arose from both directed evolution and machine learning. This was done by combining all 1.25X fold change or higher single mutants and all combinatorial double mutants with all directed evolution mutants. In addition, all combinatorial variants from Round 1 were stacked with all directed evolution mutants. The top 10 mutants were selected and assayed.

For all combinatorial predictions, EVOLVEpro was trained with N481V as a beneficial mutation (initially counted as 1.4X fold improvement), however further testing showed N481V is a neutral mutation. This reassessment does not affect any other tested substitutions, whose values remain the same (within error). Interestingly, this gives us the opportunity to assess the sensitivity of programs such as EVOLVEpro to variance.

## Results

### Zero-shot prediction of improved HpaBC variants

While a variety of models have been developed that can be used in zero-shot prediction of protein variants,^10^ and benchmarking has begun to be carried out^29–31^, *de novo* experimental validations of predictions are still relatively rare. In order to improve the catalytic efficiency of HpaBC, we explored predictions made by several models that had previously been shown to be useful for protein engineering: structure-based models such as MutCompute^11,12,18,32^ and ProteinMPNN^20^, and various releases of the popular ESM protein language model^28^.

In each instance, the algorithms were used to predict the top 60 variants based on either the monomer or (functional) tetrameric structure of the HpaB monooxygenase (HpaC contains the FAD cofactor). The overlaps within and between the different structures and methods eventually resulted in 329 predicted variants (**Figure 1**; **Supplemental Figure 1**; **Supplemental Figure 2**).

**Figure 1:**
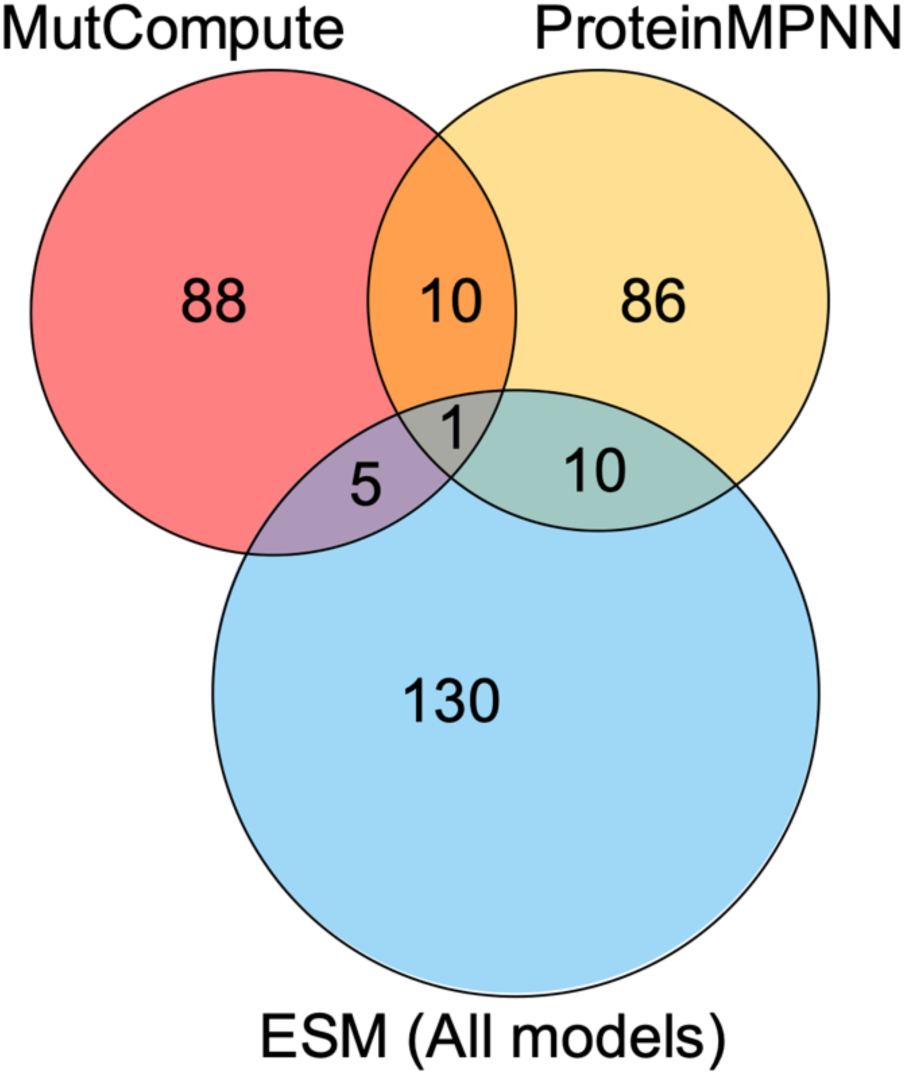
Overlap of the top predicted single-substitution variants identified by each machine learning model. Shared regions indicate mutations predicted by multiple models.

### Assaying predicted HpaBC variants

To assay these variants, we used a two-plasmid circuit, consisting of an expression vector for the enzyme and a reporter vector that we had previously developed^9^ (**Figure 2a**). The expression vector constitutively expresses HpaC and places the HpaB gene under the control of TetR via the p_Ltet0_ promoter. The reporter vector contains a circuit in which GFP production is controlled by a variant of the transcription factor DopA^9,33^ that activates transcription in the presence of L-Dopa and is known to have a large dynamic range. We^34^ and others^35^ have previously found that the use of reporter circuits to optimize production pathways for small molecules, whether by directed evolution^9^ or machine learning^15^, can lead to rapid improvements in enzyme function. In control experiments in which aTc was used to induce HpaB to different levels, the assay proved to be a reliable indicator of the amount of L-Dopa produced (**Supplemental Figure 3**).

**Figure 2:**
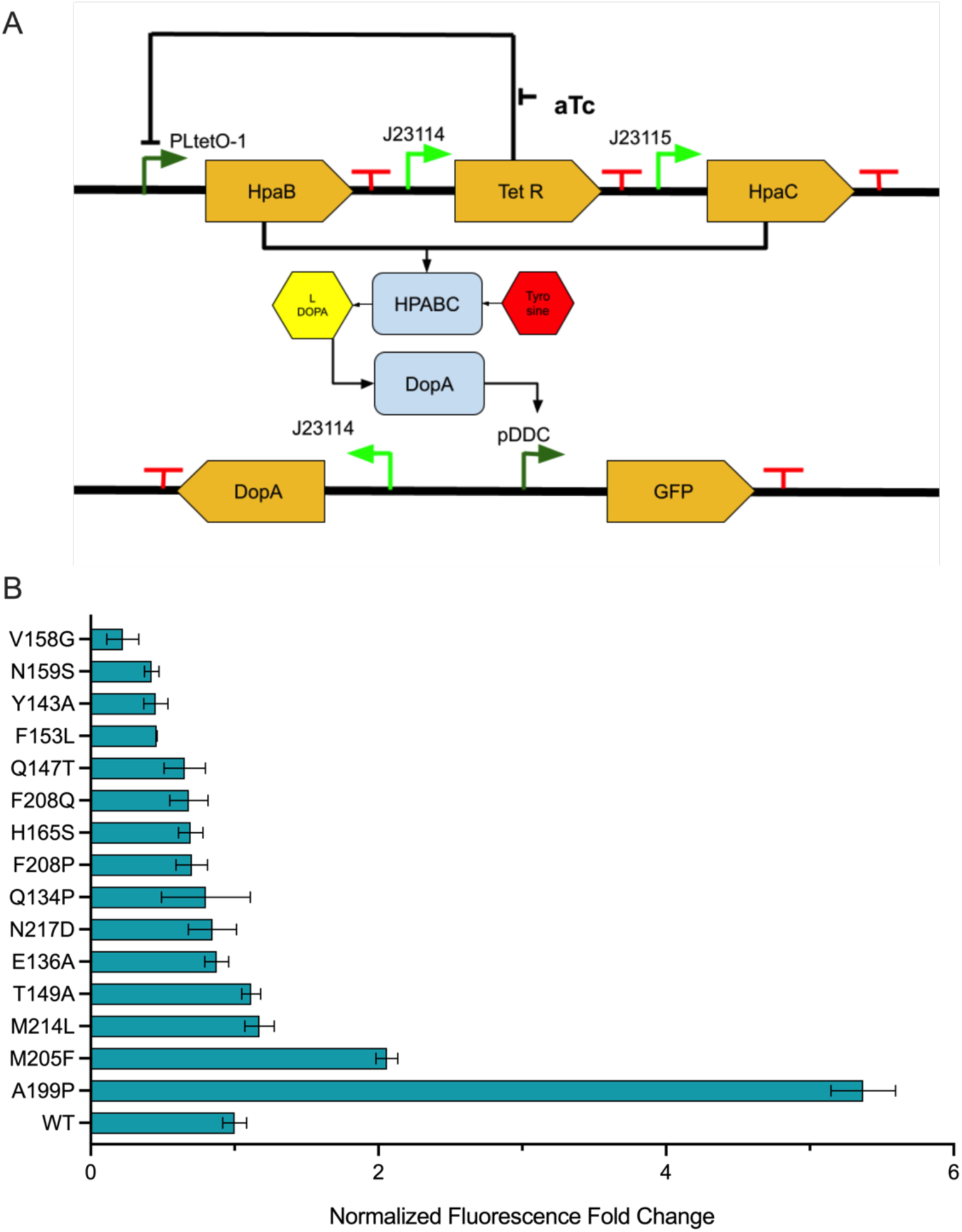
Biosensor assay for evaluating HpaBC activity and screening machine-learning-predicted variants. (A) Schematic of the L-Dopa biosensor used to quantify HpaBC activity. HpaBC catalyzes the conversion of L-tyrosine to L-Dopa, which is detected by the transcription factor DopA, activating GFP expression from the *pDDC* promoter. HpaB expression is controlled by the aTc-inducible *P*LtetO-1/TetR system. (B) Activity of machine-learning-predicted HpaBC single-substitution variants measured using the L-Dopa biosensor. Fluorescence values were normalized to cell density, background subtracted using uninduced controls and expressed as fold change relative to wild-type HpaBC measured on the same plate. Error bars represent the standard deviation of biological triplicates.

Each variant was cloned into the expression vector and assayed in triplicate following induction at 20nM aTc, a midpoint concentration that allows us to differentiate high performing variants of HpaBC. The level of fluorescence in uninduced cultures was subtracted from the level observed in induced, and this value was then normalized to the fluorescence observed in cells transformed with the wild-type HpaBC complex (**Figure 2B**). Of the 329 variants that were assayed 19 showed a change in fluorescence of 1.25x-fold or greater (**Figure 3**).

**Figure 3:**
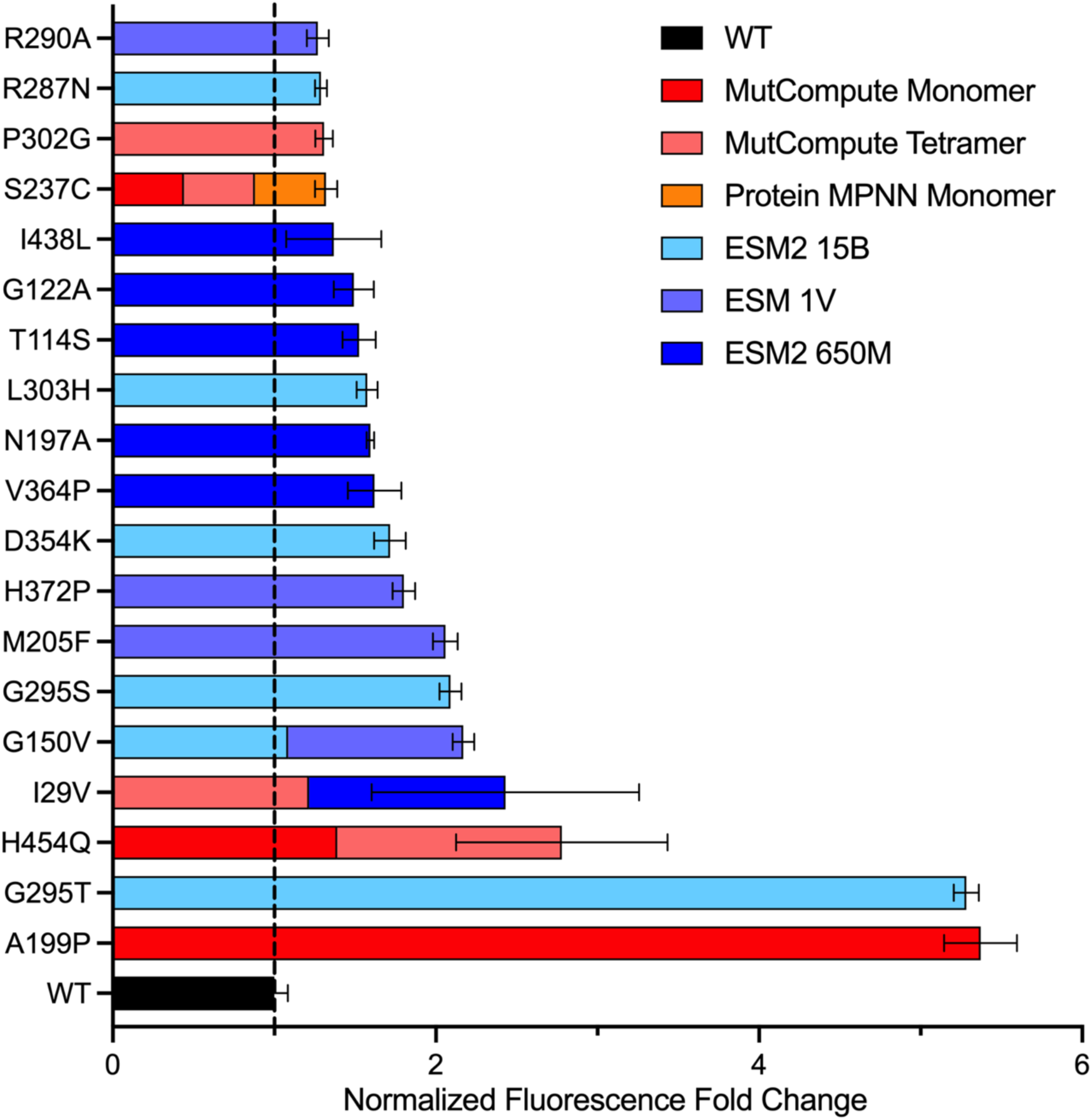
Top-performing HpaBC variants identified by machine learning. HpaBC variants exhibiting a ≥1.25-fold increase in activity relative to wild-type, as measured by the L-Dopa biosensor assay. Bar colors denote the machine learning model(s) that predicted each variant; multicolored bars indicate variants independently identified by multiple models. The dashed line indicates wild-type HpaBC activity. Error bars represent the standard deviation of biological triplicates.

At the extremes, the variant A199P (predicted by MutCompute based on the monomer input structure) showed a 5.4x-fold improvement in fluorescence output, while G295T and G295S (predicted by ESM2 15B) demonstrated 5.3x- and 2.1x-fold changes in fluorescence. It should be noted that G295T had also previously been identified via directed evolution experiments to increase L-Dopa production^9^.

### Using EVOLVEpro to Stack Single Substitutions

We sought to further improve the production of L-Dopa by combining single substitution variants. To rationalize what combinations to make we relied on the EVOLVEpro framework^23^, which trains a top layer Random Forest model to predict experimental outcome from protein language model embeddings. This trained model is then used to predict the experimental outcome of novel mutants either as combinatorial or new single substitutions. Starting from variants with 1.25x or greater changes in fluorescence predictions were generated that contained up to 5 substitutions.

In the first round of screening the top 5 predicted double, triple, quadruple, and quintuple substitutions were predicted and assayed. Many of these stacked variants yielded large fold-changes in fluorescence, indicating improved L-Dopa production (**Figure 4, Supplementary Table 3**). For example, a quintuple substitution variant, D354K_A199P_I29V_H454Q_H372P, yielded a fold-change in fluorescence of 29x, outperforming any of its constituent single substitutions garnered from both MutCompute and ESM. Interestingly, the initial two top performing single substitution variants (G295T and G295S from ESM) both performed poorly when stacked with other substitutions, despite being heavily favored by EVOLVEpro to create combinatorial variants. This indicates that the scoring mechanism of EVOLVEpro in predicting assay results lacks the fine-grained interpretation to allow for prediction of epistatic relationships between mutations.

**Figure 4:**
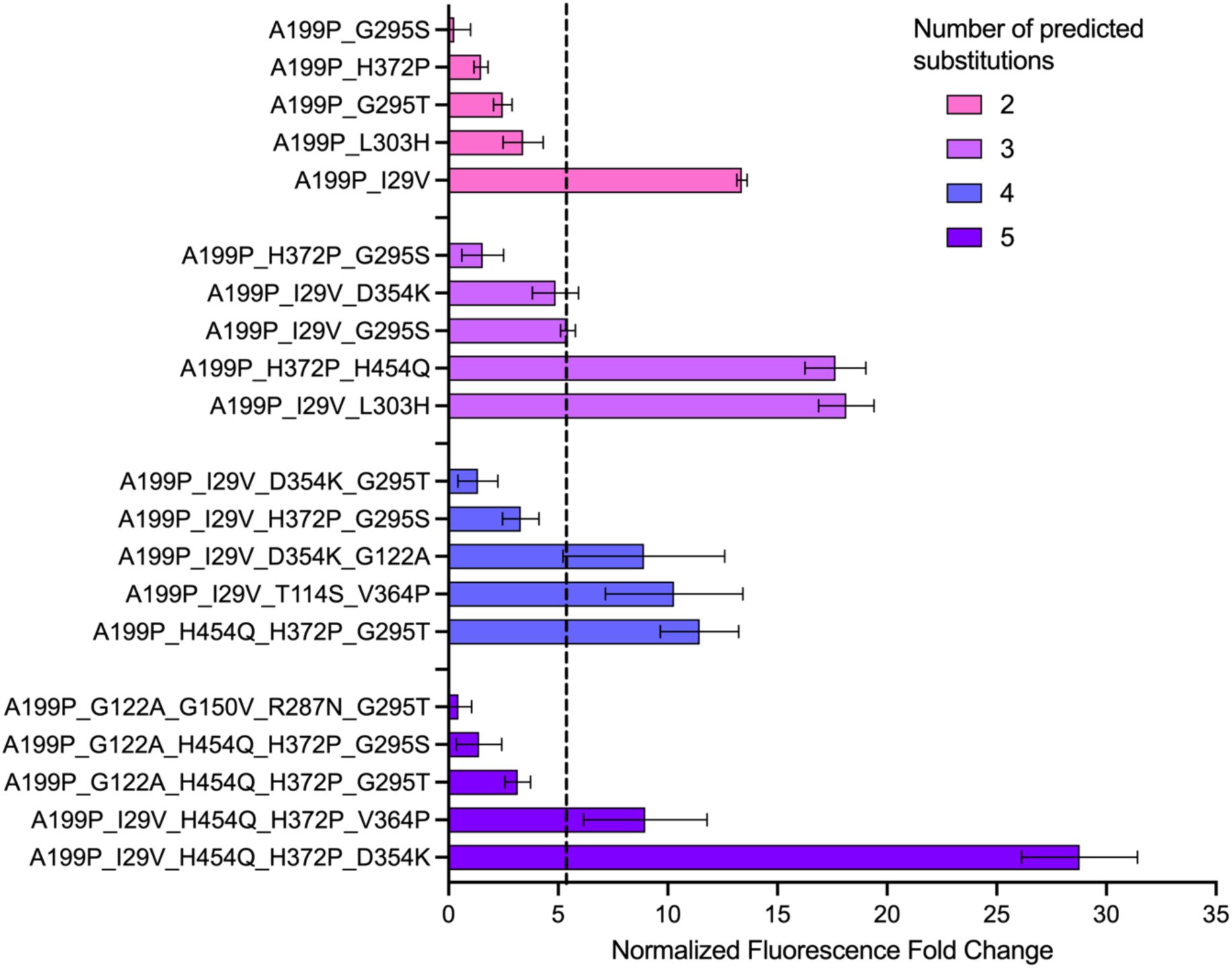
EVOLVEpro combines beneficial single substitutions to generate improved HpaBC variants. HpaBC variants containing two to five amino acid substitutions predicted by EVOLVEpro were evaluated using the L-Dopa biosensor assay. Variants are grouped by the number of substitutions. The dashed line indicates the activity of the best-performing single-substitution variant (A199P). Error bars represent the standard deviation of biological triplicates. Variant sequences are provided in Table S3 in order of EVOLVEpro rank.

### Experimental Feedback Improves EVOLVEpro Design

To determine whether incorporating experimental feedback could further improve combinatorial variant design, EVOLVEpro was retrained using the experimental results from the first round of higher order variants and then used to predict up to hextuple substitutions (**Figure 5**). In five of the six top predictions, EVOLVEpro favored an existing quintuple variant and merely added a single additional substitution (**Supplementary Table 4**), indicating that EVOLVEpro favors known successes over the exploration of entirely new mutational combinations and paths.

**Figure 5:**
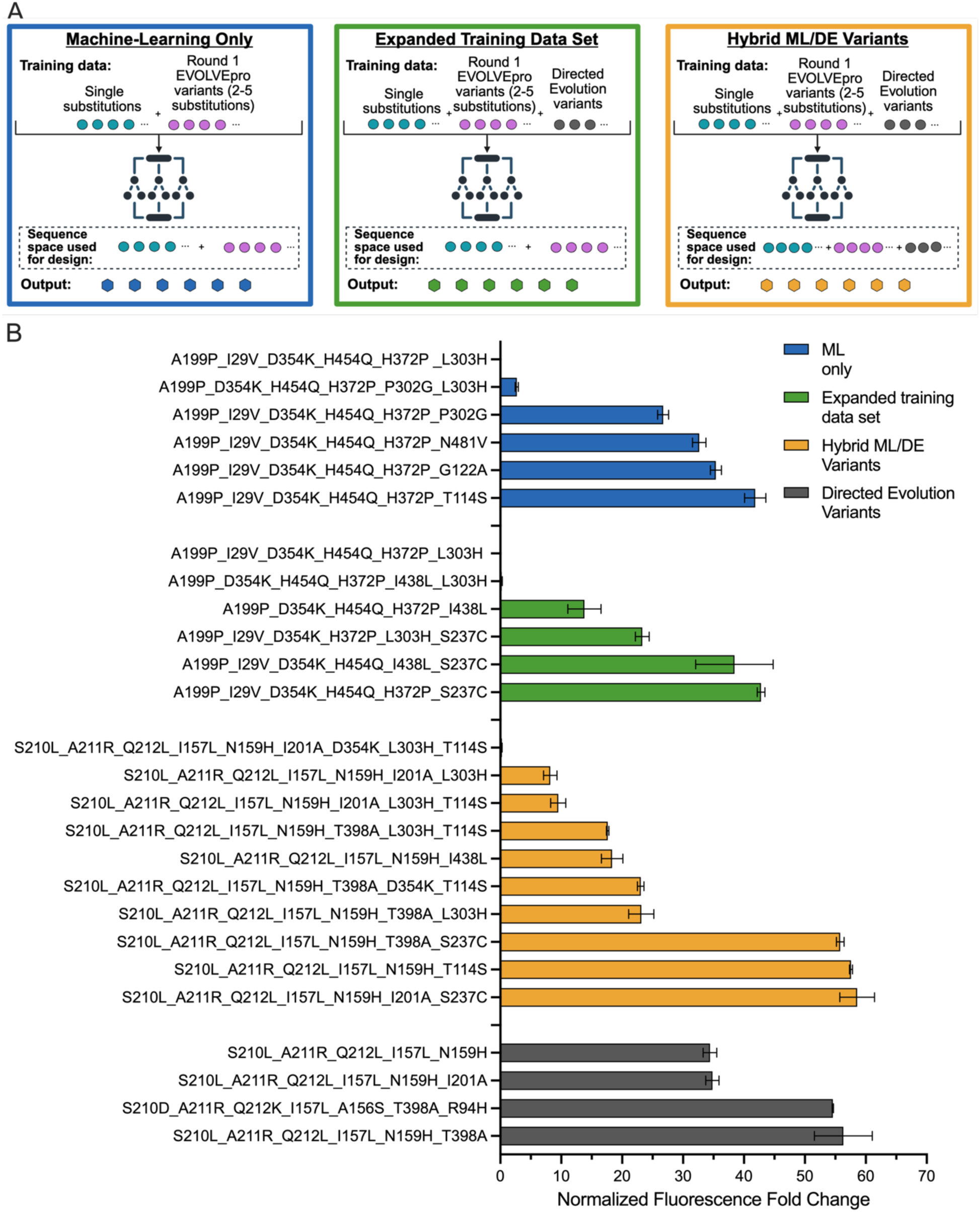
Iterative EVOLVEpro strategies for higher-order HpaBC engineering. (A) Schematic illustrating the three iterative EVOLVEpro workflows evaluated after the initial combinatorial design round: retraining with experimentally characterized EVOLVEpro variants, expanding the training dataset with directed-evolution-derived variants, and expanding the combinatorial search space to include directed-evolution backgrounds. (B) Activity of second-generation HpaBC variants predicted by each EVOLVEpro strategy, measured using the L-Dopa biosensor assay following induction with 10 nM aTc. Error bars represent the standard deviation of biological triplicates. Sequences can be found in Table S4 in order of EVOLVEpro rank.

During characterization of these second-generation variants, the fluorescence assay reached its upper detection limit when protein expression was induced with 20 nM aTc. Reducing the inducer concentration to 10 nM increased the dynamic range of the assay and enabled improved discrimination among the highest performing variants. Under these conditions, the best second-generation variant (D354K_A199P_I29V_T114S_H454Q_H372P) exhibited a 42-fold increase in fluorescence relative to the wild-type enzyme, demonstrating that iterative incorporation of experimental data enabled continued improvements in variant performance.

### Directed Evolution Data Can Expand the Training Landscape for EVOLVEpro

To evaluate how additional data from directed evolution experiments might further improve combinatorial design, EVOLVEpro was supplemented with a training set derived from a directed evolution campaign of HpaBC^9^ (**Figure 5**) and again used to suggest novel variants with up six substitutions. Incorporation of the directed evolution data led to the prediction of novel combinations of individual substitutions, with four of the six highest-ranked hextuple variants representing unique combinations, rather than single additions to previously successful quintuple variants (**Supplemental Table 4**). The newly predicted hextuple variants were synthesized and experimentally characterized. As in the previous round, one variant (D354K_S237C_A199P_I29V_H454Q_H372P) exhibited a 43-fold increase in fluorescence relative to the wild-type enzyme when assayed at 10 nM aTc.

Interestingly, one variant that was highly ranked by both the machine-learning-only and expanded training strategies (D354K_L303H_A199P_I29V_H454Q_H372P) was found to be nonfunctional, highlighting the continued challenge of predicting higher-order epistatic interactions. Although incorporating directed evolution data substantially changed EVOLVEpro’s ranking of candidate variants, it did not yield variants with greater activity than those identified using the machine-learning-derived training set alone.

### Expanding the sequence space for EVOLVEpro Combinatorial Mutants

Given the bias of EVOLVEpro to favor sequences with more substitutions, we carried out a final set of predictions that allowed up to 12-tuple substitutions to be considered, this included scoring of variants derived from directed evolution, rather than just being used as part of the training data, as above. In the end, of the top 10 variants predicted, two were hextuples, four were heptuples, three were octuples, and one was a nine-tuple. We synthesized and assayed these top 10 variants (**Figure 5, Supplemental Table 4**). This approach resulted in variants of HpaBC with the greatest increase in L-Dopa production: T114S_S210L_A211R_Q212L_I157L_N159H added a single zero shot prediction (T114S) to a previous quintuple directed evolution variant and yielded a 57-fold change in fluorescence, while S237C_S210L_A211R_Q212L_I157L_N159H_I201A added a different zero-shot prediction (S237) to a previous hextuple directed evolution variant and yielded a 58-fold change in fluorescence.

## Discussion

Engineering HpaBC to improve L-Dopa production remains an important objective for the development of sustainable biosynthetic pathways for pharmaceuticals and other catechol-containing compounds. In this study, we identified multiple HpaBC variants with substantially improved activity through an iterative machine learning-guided engineering strategy. Successive rounds of computational prediction, experimental validation, and combinatorial design produced variants with up to 58-fold higher activity than the wild-type enzyme. The best variant compared favorably with enzymes derived through directed evolution alone^9^.

This study was designed to allow comparisons between different zero-shot prediction methods (**Figure 3**). For the top 60 predicted single substitutions from each model, ESM 2 650M and ESM2 15B found 6 function enhancing substitutions (10% hit rate), while ESM 1V found 4 substitutions for (6.7% hit rate). MutCompute found 3 improved substitutions based on the predicted monomer structure and 4 improved substitutions based on the predicted oligomer structure (5% and 6.7% hit rates, respectively). Finally, ProteinMPNN found 1 substitution (1.7% hit rate) based on the predicted monomer structure, but no substitutions in its top 60 predictions using the predicted tetramer structure.

While there were roughly similar hit rates for all the algorithms tested, there was surprisingly little overlap between predicted substitutions. Of 19 variants that showed at least 1.25x fold change fluorescence or greater, I29V was predicted by both ESM2 650M and MutCompute (based on the monomer structure); G150V was predicted by both ESM1V and ESM2 15B; and S237C was predicted by both MutCompute and ProteinMPNN. The remaining 16 variants were unique to a given algorithm. Moreover, there was little correlation between the ranking of a given prediction and the activities that were experimentally observed (**Supplemental Figure 5**).

To further improve HpaBC activity, we employed EVOLVEpro to iteratively design higher-order variants from experimentally validated substitutions. In the initial round, the top five predicted double, triple, quadruple, and quintuple variants were experimentally characterized. Of these 1/5 double substitutions, 2/5 triple substitutions, 3/5 quadruple substitutions, and 2/5 quintuple substitutions outperformed the variants used in training (**Figure 4; Supplemental Figure 4**). Given a reasonable starting data set, EVOLVEpro relied on relatively few predictions to effectively identify increasingly productive combinations of beneficial substitutions, making it extremely suitable for many synthetic biology and protein engineering applications.

The experimentally characterized higher-order variants were subsequently incorporated into a second round of EVOLVEpro predictions. In general, rather than exploring entirely new combinations of substitutions, EVOLVEpro preferentially refined existing high performing variants. This was found to be consistent despite changes in training data --whether the model was provided mutants derived from ML pipelines, or when the training set was expanded to include directed evolution data.

Despite seemingly conservative predictions in the number of substitutions selected, EVOLVEpro was found to be very successful in guiding selection of improved HpaBC variants. The best quintuple variant from the initial training was 29-fold better than wild-type; the best hextuple built on top of this quintuple was 42-fold better. When directed evolution data was introduced, an evolved quintuple variant became a hextuple variant that was 43-fold better than wild-type, while an evolved hextuple variant became a heptuple variants was 58-fold better. In each instance, only a relatively small number of variants (6 to 10) were built and tested.

The success of EVOLVEpro appears to be from successful identification of additive or synergistic mutation effects. However, this was not always the case: when the suggestion substitution G295T was added to multi-mutant backgrounds, there was a general decrease in activity. However, enough constructive combinations existed resulting in highly active enzymes. The fitness landscapes identified in both machine learning and directed evolution methods were not incongruent, as EVOLVEPro was able to successfully combine zero-shot mutations with those that derived from directed evolution to for the greatest increase in enzyme activity.

Deep learning models have demonstrated some success in identifying beneficial single mutations, with different methods often yielding functionally similar outcomes. However, the question remains of how to engineer across multi-substitution landscapes to identify highly optimized enzymes. Directed evolution naturally explores mutational trajectories and can readily traverse landscapes that include epistatic interactions. Thus, if a high-throughput assay is available, it remains the best means of exploring high dimensional fitness landscapes. Our work suggests that models trained solely on single mutant data have limited ability to infer epistatic interactions and avoid deleterious pairings, but new algorithms (e.g., MULTIevolve^36^) that actively interpret rounds of screening data should prove increasingly useful for predicting higher dimensional fitness landscapes.

Given that other studies^9^ have reliably shown that L-Dopa titers follow enzymatic activities, as measured by the DopA reporter circuit, it is likely that the enzymes, circuits, and strains generated herein may prove useful for the eventual manufacture of this high-value intermediate, and could be combined with other downstream synthesis steps to produce diverse alkaloids on demand^37^.

## Acknowledgements and Funding

This material is based on research sponsored by the Naval Surface Warfare Center Crane Division under agreement number FA8650-21-2-5028. The U.S. Government is authorized to reproduce and distribute reprints for Governmental purposes notwithstanding any copyright notation thereon.

The views and conclusions contained herein are those of the authors and should not be interpreted as necessarily representing the official policies or endorsements, either expressed or implied, of Navy or the U.S. Government.

This project was made possible with the support of The Bioindustrial Manufacturing and Design Ecosystem (BioMADE); the content expressed herein is that of the authors and does not necessarily reflect the views of BioMADE.

## Author Contributions

D.G. and A.D.E.: Conceptualization; D.G.: investigation and formal analysis; D.G., I.M.H., and A.D.E.: writing original draft; D.G., I.M.H., A.F., and A.D.E.: writing review; D.G. and I.M.H.: visualizations; A.D.E.: supervision; A.D.E.: funding acquisition. The manuscript was written through contributions of all authors. All authors have given approval to the final version of the manuscript.

