## Supplemental Figures for "Combining Machine Learning and Directed Evolution for Optimization of a Monooxygenase"

**ORCIDs:** D.G., 0009-0002-4484-9690; I.M.H., 0009-0005-6028-2809; A.F., 0000-0002-4476-1026; A.D.E., 0000-0001-6246-5338

This file contains:
5 Supplementary Figures
4 Supplementary Tables

**Supplemental Figures:**

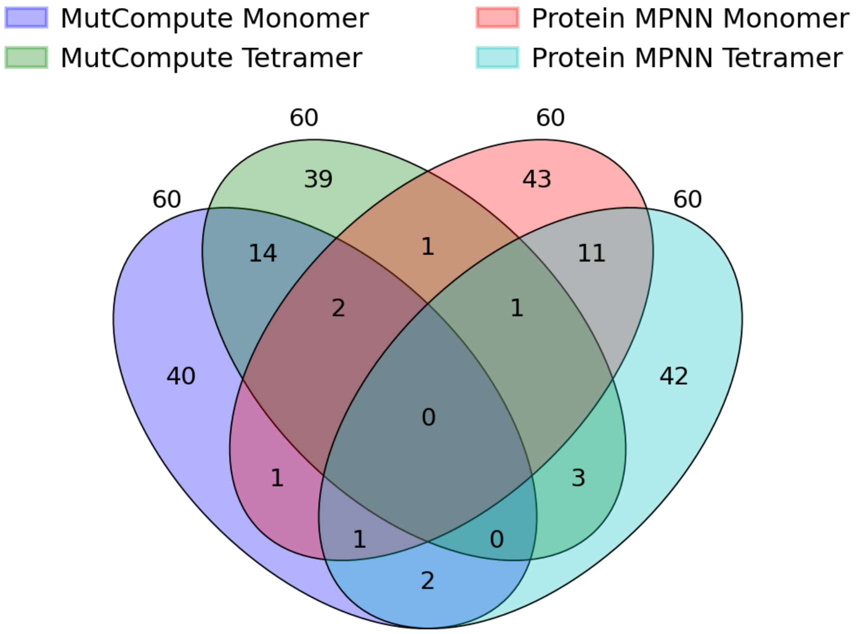

**Supplemental Fig 1:** The overlap of top 60 predictions from MutCompute and ProteinMPNN model using both a monomer HpaB as the input structure and the tetramer input structure.

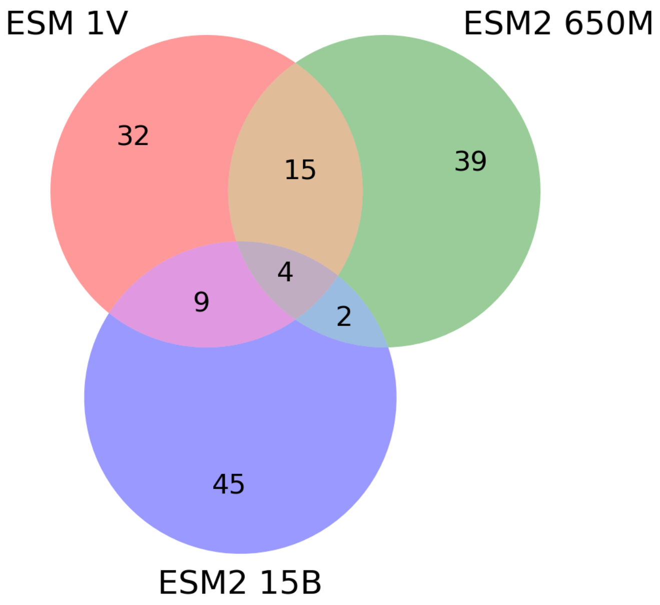

**Supplemental Fig 2:** The overlap in top 60 single mutant predictions between ESM1V, ESM2 650M, and ESM2 15B.

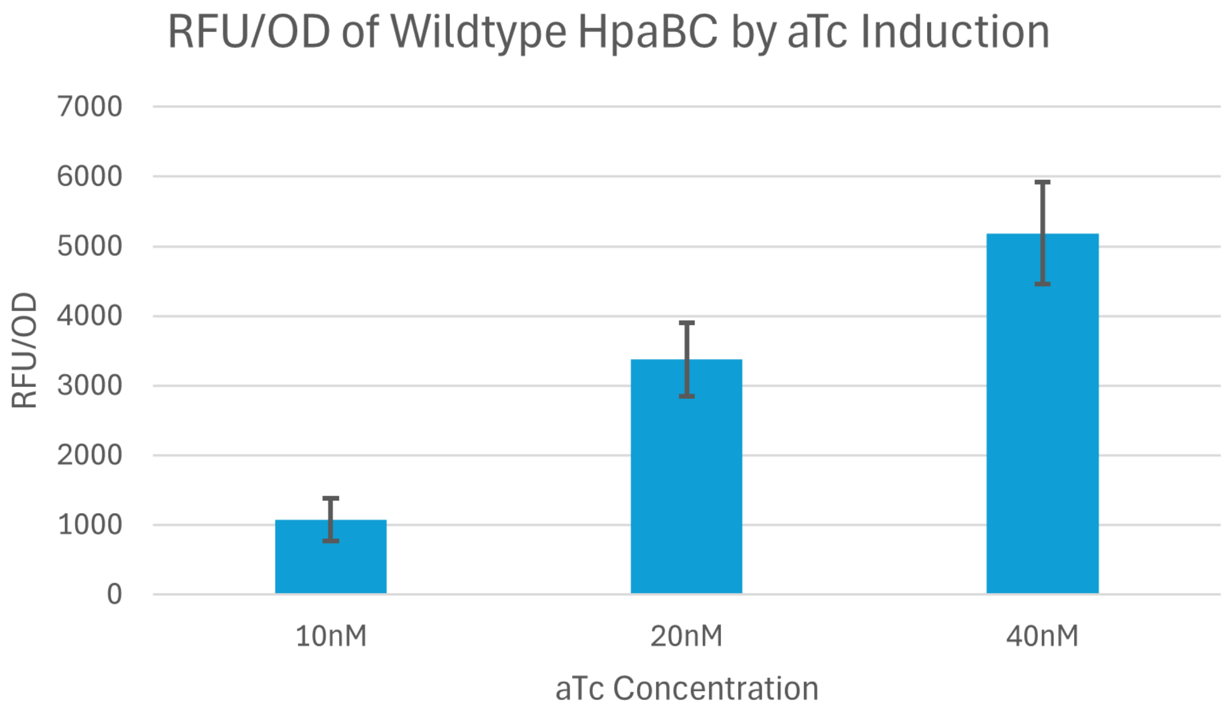

**Supplemental Fig 3:** Fluorescence of the wildtype HpaBC when induced at varying concentrations of aTc**,** higher concentrations of aTc resulted in greater HpaBC production and thus L-Dopa production as measured by the levels of GFP.

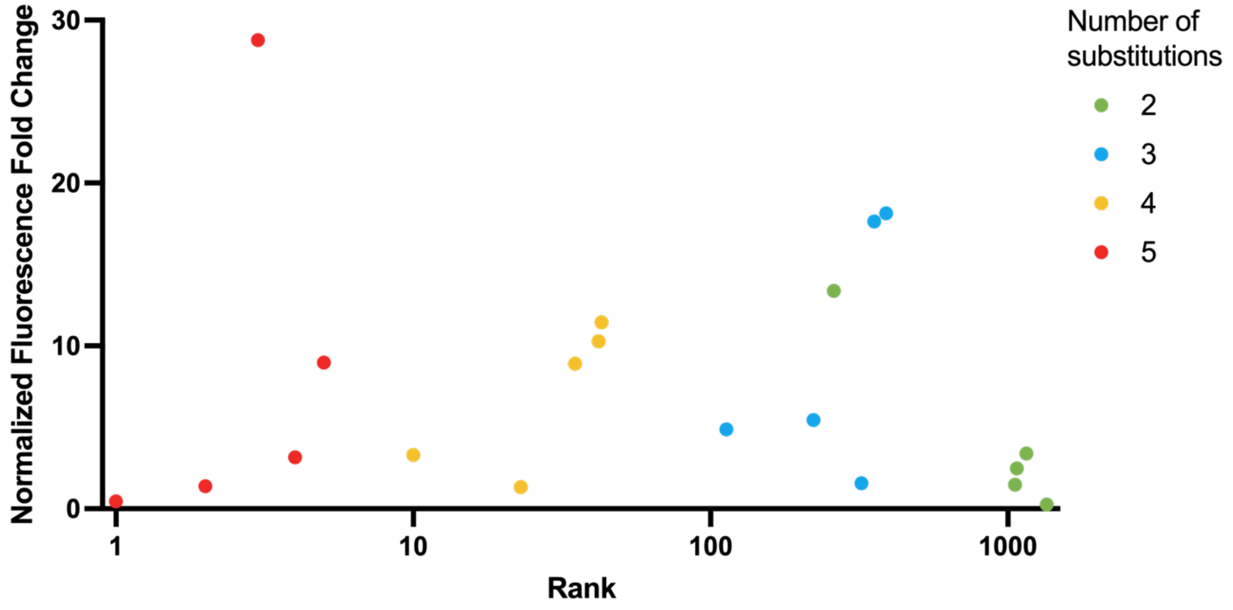

**Supplemental Fig 4:** EVOLVEpro rankings of all possible variants in round 1 versus normalized fold change in fluorescence.

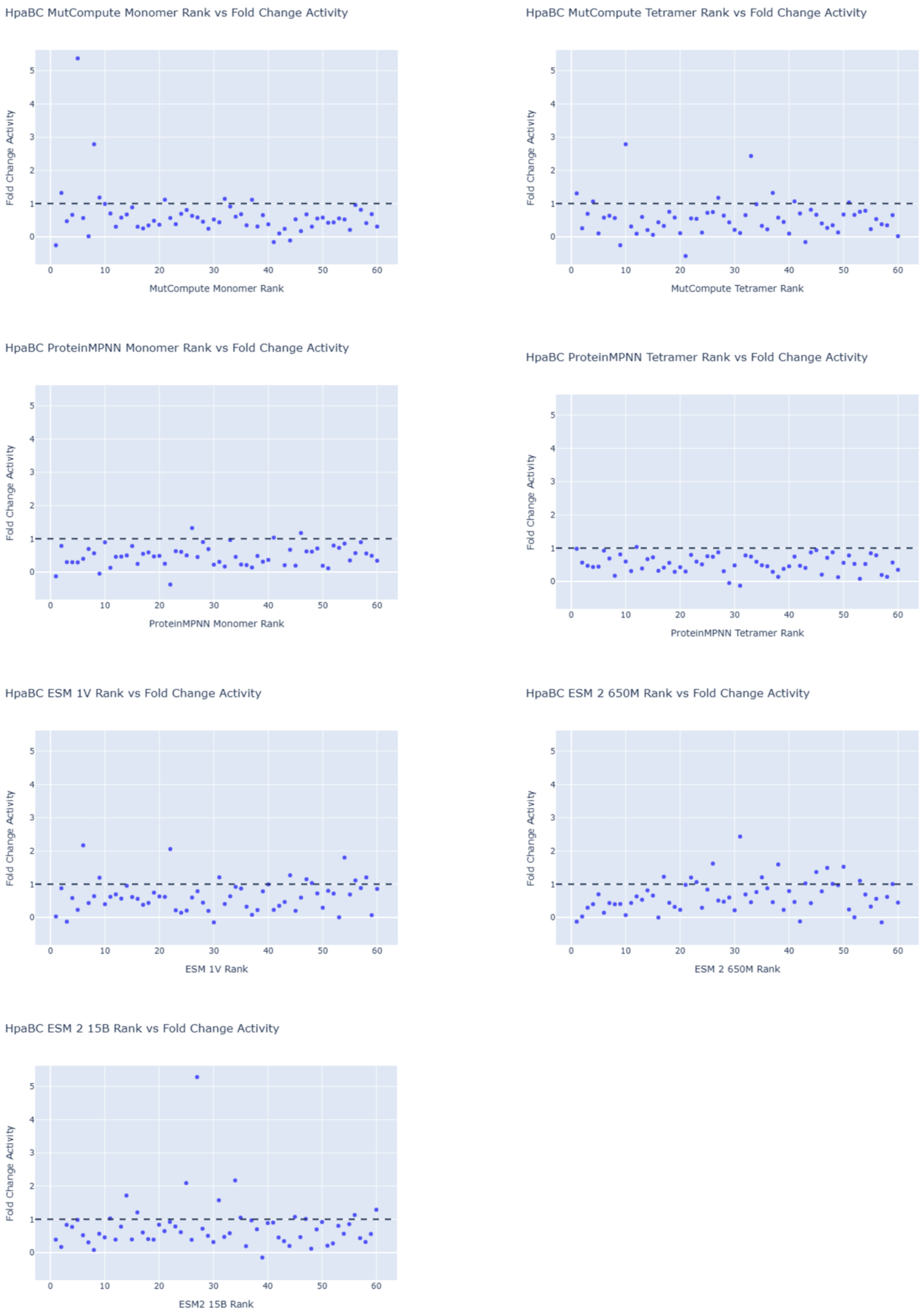

**Supplemental Fig 5**: ML rank generated by each zero-shot model versus normalized fold change in fluorescence.

**Supplemental Tables:**

Table S1: Plasmid backbones used in this study.  BsaI cut sites shown in upper case letters.

| | |
|---|---|
| WT HpaBC Plasmid | ttgagatcctttttttctgcgcgtaatctgctgcttgcaaacaaaaaaaccaccgctaccagcggtggtttgtttgccg<br>gatcaagagctaccaactctttttccgaaggtaactggcttcagcagagcgcagataccaaatactgtccttctagt<br>gtagccgtagttaggccaccacttcaagaactctgtagcaccgcctacatacctcgctctgctaatcctgttaccag<br>tggctgctgccagtggcgataagtcgtgtcttaccgggttggactcaagacgatagttaccggataaggcgcagc<br>ggtcgggctgaacggggggttcgtgcacacagcccagcttggagcgaacgacctacaccgaactgagatacct<br>acagcgtgagctatgagaaagcgccacgcttcccgaagggagaaaggcggacaggtatccggtaagcggca<br>gggtcggaacaggagagcgcacgagggagcttccagggggaaacgcctggtatctttatagtcctgtcgggttt<br>cgccacctctgacttgagcgtcgatttttgtgatgctcgtcaggggggcggagcctatggaaaaacgccagcaac<br>gcggcctttttacggttcctggccttttgctggccttttgctcacatgttctttcctgcgttatcccctgattctgtggata<br>accgtattaccgcctttgagtgagctgataccgctcgccgcagccgaacgactgagcgcagcgagtcagtgagc<br>gaggaagcggaagagcgcctgatgcggtattttctccttacgcatctgtgcggtatttcacaccgcaatggtgcac<br>tctcagtacaatctgctctgatgccgcatagttaagccagtatacactccgctatcgctacgtgactgggtcatggct<br>gcgccccgacacccgccaacacccgctgacgcgccctgacgggcttgtctgctcccggcatccgcttacagac<br>aagctgtgaccatctccgggagctgcatgtgtcagaggttttcaccgtcatcaccgaaacgcgcgaggcagctgc<br>ggtaaagctcatcagcgtggtcgtgaagcgattcacagatgtctgcctgttcatccgcgtccagctcgttgagtttct<br>ccagaagcgttaatgtctggcttctgataaagcgggccatgttaagggcggttttttcctgtttggtcactgatgcctc<br>cgtgtaagggggatttctgttcatgggggtaatgataccgatgaaacgagagaggatgctcacgatacgggttact<br>gatgatgaacgtatatatgagtaaacttggtctgacagaaagcaagctgataaaccgatacaattaaaggctcctttt<br>ggagcctttttttttggagattttcaacatgaaaaaattattattgcggccgcccctgaattcgcatctagactgaactg<br>gccgataattgcagacgaacggcgatacagaccctaatttcacatcatatgacactaatccctatcagtgatagag<br>attgacatccctatcagtgatagagatactgagcacatcagcaggacgcactgaccggagagctgtcaccggat<br>gtgctttccggtctgatgagtccgtgaggacgaaacagcctctacaaataattttgtttaagactttaaagtccaatat<br>gaaaccagaagatttccgcgccagtacccaacgtccgttcaccggggaagagtatctgaaaagcctgcaggatg<br>gtcgcgagatctatatctatggcgagcgagtgaaagacgtcactactcatccggcatttcgtaatgcggctgcgtct<br>gttgcccaactgtacgacgcgctacacaaaccggagatgcaggactctctgtgctggaacaccgacaccggca<br>gcggcggctatacccataaattcttccgcgtggcgaaaagtgccgacgacctgcgccagcaacgcgatgccatc<br>gctgagtggtcacgcctgagctatggctggatgggccgtaccccagactacaaagctgctttcggttgtgcactg<br>ggcgcgaatccgggcttttacggtcagttcgagcagaacgcccgtaactggtacacccgtattcaggaaactggc<br>ctctactttaaccacgcgattgttaacccaccgatcgatcgtcatttgccgaccgataaagtaaaagacgtttacatc<br>aagctggaaaaagagactgacgccgggattatcgtcagcggtgcgaaagtggttgccaccaactcggcgctga<br>ctcactacaacatgattggcttcggctcggcacaagtaatgggcgaaaacccggacttcgcactgatgttcgttgc<br>gccaatggatgccgatggcgtcaaattaatctcccgcgcctcttatgagatggtcgcgggtgctaccggctcacc<br>gtatgactacccgctctccagccgcttcgatgagaacgatgcgattctggtgatggataacgtgctgatcccatgg<br>gaaaacgtgctgatctaccgcgattttgatcgctgccgtcgctggacgatggaaggcggtttcgcccgtatgtatc<br>cgctgcaagcctgtgtgcgcctggcagtgaaactcgacttcattacggcactgctgaaaaaatcactcgaatgtac<br>cggcaccctggagttccgtggtgtgcaggccgatctcggtgaagtggtggcgtggcgcaacaccttctgggcatt<br>gagtgactcgatgtgttctgaagcgacgccgtgggtcaacggggcttatttaccggatcatgccgcactgcaaac<br>ctatcgcgtactggcaccaatggcctacgcgaagatcaaaaacattatcgaacgcaacgttaccagtggcctgat<br>ctatctcccttccagtgcccgtgacctgaacaatccgcagatcgaccagtatctggcgaagtatgtgcgcggttcg<br>aacggtatggatcacgtccagcgcatcaagatcctcaaactgatgtgggatgccattggcagcgagtttggtggtc<br>gtcacgaactgtatgaaatcaactactctggtagccaggatgagattcgcctgcagtgtctgcgccaggcacaaa<br>gctccggcaatatggacaagatgatggcgatggttgatcgctgcctgtcggaatacgaccagaacggctggact |

| | |
|---|---|
| | gtgccgcacctgcacaacaacgacgatatcaacatgctggataagctgctgaaataaggatccttcgtcggtcag<br>tttcacctgatttacgtaaaaacccgcttcggcgggtttttgcttttggaggggcagaaagatgaatgactgtcggcc<br>attccgggtagcataacccccttggggcctctaaacgggtcttgaggggttttttgtcatgtgaggctgccaaaccag<br>atgtcaacacagctacaacgtttatggctagctcagtcctaggtacaatgctagcggagagccccatagggtggt<br>gtgtaccacccctgatgagtccaaaaggacgaaatggggcctctacaaataattttgtttaacgtaactacggtacc<br>tatgtctcgtttagataaaagtaaagtgattaacagcgcattagagctgcttaatgaggtcggaatcgaaggtttaac<br>aacccgtaaactcgcccagaagctaggtgtagagcagcctacattgtattggcatgtaaaaaataagcgggctttg<br>ctcgacgccttagccattgagatgttagataggcaccatactcacttttgccctttagaaggggaaagctggcaag<br>atttttacgtaataacgctaaaagttttagatgtgctttactaagtcatcgcgatggagcaaaagtacatttaggtaca<br>cggcctacagaaaaacagtatgaaactctcgaaaatcaattagcctttttatgccaacaaggtttttcactagagaat<br>gcattatatgcactcagcgcagtggggcattttactttaggttgcgtattggaagatcaagagcatcaagtcgctaa<br>agaagaaagggaaacacctactactgatagtatgccgccattattacgacaagctatcgaattatttgatcaccaag<br>gtgcagagccagccttcttattcggccttgaattgatcatatgcggattagaaaaacaacttaaatgtgaaagtgggt<br>cttaaatcctaactcgaggggaactgccaggcatcaaataaaacgaaaggctcagtcggaagactgggcctttcg<br>ttttatctgttgtttgtcggtgaacgctctcctggctggatgcacacactggcttaagatgacaacgtttatagctagct<br>cagcccttggtacaatgctagcggagagtcataagtctgggctaagcccactgatgagtcgctgaaatgcgacga<br>aacttatgacctctacaaataattttgtttaatgaaccacgaggccttatgcaattagatgaacaacgcctgcgctttc<br>gtgacgcaatggccagcctgtcggcagcggtaaatattatcaccaccgagggcgacgccggacaatgcgggat<br>tacggcaacggccgtctgctcggtcacggatacaccaccatcgctgatggtgtgcattaacgccaacagtgcgat<br>gaacccggtttttcagggcaacggtaagttgtgcgtcaacgtcctcaaccatgagcaggaactgatggcacgcca<br>cttcgcgggcatgacaggcatggcgatggaagagcgttttagcctctcatgctggcaaaaaggtccgctggcgc<br>agccggtgctaaaaggttcgctggccagtcttgaaggtgagatccgcgatgtgcaggcaattggcacacatctgg<br>tgtatctggtggagattaaaaacatcatcctcagtgcagaaggtcacggacttatctactttaaacgccgtttccatc<br>cggtgatgctggaaatggaagctgcgatttaaggatcctaactcgaggggaactgccaggcatcaaataaaacg<br>aaaggctcagtcgaaagactgggcctttcgttttatctgttgtttgtcggtgaacgctctcctggctgagcagttaca<br>gagatgttacgaaccacgcggccgcgattatcaaaaaggatcttcacctagatccttttaaattaaaaatgaagtttt<br>aaatcaatctaaagtatatatgagtaaacttggtctgacagttaccaatgcttaatcagtgaggcacctatctcagcg<br>atctgtctatttcgttcatccatagttgcctgactccccgtcgtgtagataactacgatacgggagggcttaccatctg<br>gccccagtgctgcaatgataccgcgggacccacgctcaccggctccagatttatcagcaataaaccagccagcc<br>ggaagggccgagcgcagaagtggtcctgcaactttatccgcctccatccagtctattaattgttgccgggaagcta<br>gagtaagtagttcgccagttaatagtttgcgcaacgttgttgccattgctacaggcatcgtggtgtcacgctcgtcgt<br>ttggtatggcttcattcagctccggttcccaacgatcaaggcgagttacatgatcccccatgttgtgcaaaaaagcg<br>gttagctccttcggtcctccgatcgttgtcagaagtaagttggccgcagtgttatcactcatggttatggcagcactg<br>cataattctcttactgtcatgccatccgtaagatgcttttctgtgactggtgagtactcaaccaagtcattctgagaata<br>gtgtatgcggcgaccgagttgctcttgcccggcgtcaatacgggataataccgcgccacatagcagaactttaaa<br>agtgctcatcattggaaaacgttcttcggggcgaaaactctcaaggatcttaccgctgttgagatccagttcgatgta<br>acccactcgtgcacccaactgatcttcagcatcttttactttcaccagcgtttctgggtgagcaaaaacaggaaggc<br>aaaatgccgcaaaaaagggaataagggcgacacggaaatgttgaatactcatactcttcctttttcaatattattgaa<br>gcatttatcagggttattgtctcatgagcggatacatatttgaatgtatttagaaaaataaacaaataggggttccgcg<br>cacatttccccgaaaagtgccacctgggaactatatccggattggcgaatgggtcatgaccaaaatcccttaacgt<br>gagttttcgttccactgagcgtcagaccccgtagaaaagatcaaaggatcttc |
| L-Dopa Reporter Vector | acagtccagcttggagcgaactgcctacccggaactgagtgtcaggcgtggaatgagacaaacgcggccataa<br>cagcggaatgacaccggtaaaccgaaaggcaggaacaggagagcgcacgagggagccgccagggggaaa<br>cgcctggtatctttatagtcctgtcgggtttcgccaccactgatttgagcgtcagatttcgtgatgcttgtcagggggg<br>gcggagcctatggaaaaacggctttgccgcggccctctcacttccctgttaagtatcttcctggcatcttccaggaa<br>atctccgccccgttcgtaagccatttccgctcgccgcagtcgaacgaccgagcgtagcgagtcagtgagcgagg |

| | |
|---|---|
| | aagcggaatatatcctgtatcacatattctgctgacgcaccggtgcagcctttttctcctgccacatgaagcacttca<br>ctgacaccctcatcagtgccaacatagtaagccagtatacactccgctagcgctgatgtccggcggtgcttttgcc<br>gttacgcaccaccccgtcagtagctgaacaggagggacagctgatagaaacagaagccactggagcacctcaa<br>aaacaccatcatacactaaatcagtaagttggcagcatcacccgacgcactttgcgccgaataaatacctgtgacg<br>gaagatcacttcgcagaataaataaatcctggtgtccctgttgataccgggaagccctgggccaacttttggcgaa<br>aatgagacgttgatcggcacgtaagaggttccaactttcaccataatgaaataagatcactaccgggcgtattttttg<br>agttatcgagattttcaggagctaaggaagctaaaatggagaaaaaaatcactggatataccaccgttgatatatcc<br>caatggcatcgtaaagaacattttgaggcatttcagtcagttgctcaatgtacctataaccagaccgttcagctggat<br>attacggcctttttaaagaccgtaaagaaaaataagcacaagttttatccggcctttattcacattcttgcccgcctgat<br>gaatgctcatccggagttccgtatggcaatgaaagacggtgagctggtgatatgggatagtgttcacccttgttaca<br>ccgttttccatgagcaaactgaaacgttttcatcgctctggagtgaataccacgacgatttccggcagtttctacaca<br>tatattcgcaagatgtggcgtgttacggtgaaaacctggcctatttccctaaagggtttattgagaatatgtttttcgtct<br>cagccaatccctgggtgagtttcaccagttttgatttaaacgtggccaatatggacaacttcttcgcccccgttttcac<br>tatgggcaaatattatacgcaaggcgacaaggtgctgatgccgctggcgattcaggttcatcatgccgtttgtgatg<br>gcttccatgtcggcagaatgcttaatgaattacaacagtactgcgatgagtggcagggcggggcgtaatttttttaa<br>ggcagttattggtgcccttaaacgcctggttgctacgcctgaataagtgataataagcggatgaatggcagaaattc<br>gaaagcaaattcgacccggtcgtcggttcagggcagggtcgttaaatagccgcttatgtctattgctggtttaccgg<br>tttattgactaccggaagcagtgtgaccgtgtgcttctcaaatgcctgaggtttcagcaaaaaacccctcaagaccc<br>gtttagaggccccaaggggttatgctagttattgctcagcggtggcagcagcctaggttaattaagctgcgctagta<br>gacgagtccatgtgctggcgttcaaatttcgcagcagcggtttctttaccagactcgagttatttgtacagttcgtcca<br>tgtcgtgggtgatacccgccgcggtaacgaattccagcagaaccatgtggtcacgtttttcgttcgggtctttagac<br>agcgcagactgggtagacaggtagtggttgttcggcagcagaaccggaccgtcaccgatcggggtgttctgctg<br>gtagtggtccgccagctgaacagaaccgtcttcgatgttgtgacggattttgaagttaactttgataccgtttttctgtt<br>tgtccgccatgatgtaaacgttgtgagagttgtagttgtattccagtttgtgacccaggatgttaccgtcttctttgaag<br>tcgatacctttcagaacgatacggttaaccagggtgtcaccttcgaatttaacttccgcacgggttttgtagtaaccgt<br>cgtcttcgaagaagatggtacgttcctgaacgtaaccttccggcatcgcagatttggagaagtcgtgctgtttcatgt<br>ggtccgggtaacgcgcgaagcactgcagaccgtacgccagggtggtaaccagggtcggccacggaaccggc<br>agtttaccggtggtgcagatgaatttcagggtcagtttaccgtaggtcgcgtcaccttcaccttcaccagaaacaga<br>gaatttgtgaccgttaacgtcaccgtccagttcaaccaggatcggaacaacaccggtgaacagttcttcacctttag<br>acatatgtacgtacctccttagggttatccattgcctgtgctggccctttcgcgggtaaacccgcgcctacagccgc<br>gaagaagccgcacaatcaacacaatgccccagtctaggcacctatccccagccgaataacctcgcttaccgcat<br>agctgctatggagtcgcataagggagagcgtcgagatcccggacaccatcgaatggcgcaaaacctttcgcggt<br>atggcatgatagcgcccggaagagagtcaattcagggtggtgaatgcgtatcacgaggccctttcgtcttcacctc<br>gagatttatggctagctcagtcctaggtacaatgctagcgaaataattttgtttaactttattaaagaggagaaattaa<br>ctatgaatccgactttcgcaagcctgtccctggcgcacctgcgcactctggatcacctgctgcaactgaaaaacct<br>gagccacgcagcagaacgcctgggtgtatcccagtctgccctgtctcgtcagctggcccatctgcgtgaggcgtt<br>cgatgatccgctgctggtccgtcaaggccgtggttatgtactgtctgaacacgcagaggccctggtcgaaccgct<br>gcgtcaggttctggaggagctgcacgctctgcgccagcctgctatcttcgatccggcccgctgcgaacgtcgcttt<br>tgcctggcagcgagcgattacgtagcagagcatatgctgccgctgctggttgcggcactggagcgtgaagctcc<br>aggtgtttctctggagtaccgtacctggcaggcgggccaatatgctctgctggcttctggcgagatcgacctggcc<br>accaccatgttcaccgagtccccgccgaacctgcacggtcgtctgctgggtgaagatcgcgctgtatgtctgatg<br>cgtcaggatcacccgctggctgcacaggcagcgctgtctcaggcagactacctggcgtacaaacacgtgcgtat<br>cagcggtggtggtgataaagactctttcatcgatcgccatctgcgcgcccagggcctgcagcgtcgcgtgtccct<br>ggaagtaccattcttctgcgcgaccgtacaggtgatcgcgtcttctcaggctgttgccacggtgccagagcatatt<br>gctcgccagctgtcccgcctgcacgatctggcgtggcgcccgctgggcttcattgaccacagccagcgttactg<br>ggtggtttggcaccagcgcctgcaggcgtctgctgagcaccgttggctgcgtaaccgtgttttcgaactgtggcgt |

| | |
|---|---|
| | cagtctcagttcggtgtacagggtggccatgcaggttctccgtaagcgcaacgcaattaatgtaagttagctcactc<br>attaggcaccgggatctcgaccgatgcccttgagagccttcaacccagtcagctccttccggtgggcgcggggc<br>atgactaacatgagaattacaacttatatcgtatggggctgacttcaggtgctacatttgaagagataaattgcactg<br>aaatctagaaatattttatctgattaataagatgatcttcttgagatcgttttggtctgcgcgtaatctcttgctctgaaaa<br>cgaaaaaaccgccttgcagggcggtttttcgaaggttctctgagctaccaactctttgaaccgaggtaactggcttg<br>gaggagcgcagtcaccaaaacttgtcctttcagtttagccttaaccggcgcatgacttcaagactaactcctctaaa<br>tcaattaccagtggctgctgccagtggtgcttttgcatgtctttccgggttggactcaagacgatagttaccggataa<br>ggcgcagcggtcggactgaacggggggttcgtgcat |
| HpaBC<br>Golden<br>Gate<br>Backbone 1 | ttgagatcctttttttctgcgcgtaatctgctgcttgcaaacaaaaaaaccaccgctaccagcggtggtttgtttgccg<br>gatcaagagctaccaactctttttccgaaggtaactggcttcagcagagcgcagataccaaatactgtccttctagt<br>gtagccgtagttaggccaccacttcaagaactctgtagcaccgcctacatacctcgctctgctaatcctgttaccag<br>tggctgctgccagtggcgataagtcgtgtcttaccgggttggactcaagacgatagttaccggataaggcgcagc<br>ggtcgggctgaacggggggttcgtgcacacagcccagcttggagcgaacgacctacaccgaactgagatacct<br>acagcgtgagctatgagaaagcgccacgcttcccgaagggagaaaggcggacaggtatccggtaagcggca<br>gggtcggaacaggagagcgcacgagggagcttccagggggaaacgcctggtatctttatagtcctgtcgggttt<br>cgccacctctgacttgagcgtcgatttttgtgatgctcgtcaggggggcggagcctatggaaaaacgccagcaac<br>gcggcctttttacggttcctggccttttgctggccttttgctcacatgttctttcctgcgttatcccctgattctgtggata<br>accgtattaccgcctttgagtgagctgataccgctcgccgcagccgaacgactgagcgcagcgagtcagtgagc<br>gaggaagcggaagagcgcctgatgcggtattttctccttacgcatctgtgcggtatttcacaccgcaatggtgcac<br>tctcagtacaatctgctctgatgccgcatagttaagccagtatacactccgctatcgctacgtgactgggtcatggct<br>gcgccccgacacccgccaacacccgctgacgcgccctgacgggcttgtctgctcccggcatccgcttacagac<br>aagctgtgaccatctccgggagctgcatgtgtcagaggttttcaccgtcatcaccgaaacgcgcgaggcagctgc<br>ggtaaagctcatcagcgtggtcgtgaagcgattcacagatgtctgcctgttcatccgcgtccagctcgttgagtttct<br>ccagaagcgttaatgtctggcttctgataaagcgggccatgttaagggcggttttttcctgtttggtcactgatgcctc<br>cgtgtaagggggatttctgttcatgggggtaatgataccgatgaaacgagagaggatgctcacgatacgggttact<br>gatgatgaacgtatatatgagtaaacttggtctgacagaaagcaagctgataaaccgatacaattaaaggctcctttt<br>ggagcctttttttttggagattttcaacatgaaaaaattattattgcggccgcccctgaattcgcatctagactgaactg<br>gccgataattgcagacgaacggcgatacagaccctaatttcacatcatatgacactaatccctatcagtgatagag<br>attgacatccctatcagtgatagagatactgagcacatcagcaggacgcactgaccggagagctgtcaccggat<br>gtgctttccggtctgatgagtccgtgaggacgaaacagcctctacaaataattttgtttaagactttaaagtccaatA<br>GAGACCatgaaaccagaagatttccgcgccagtacccaacgtccgttcaccggggaagagtatctgaaaa<br>gcctgcaggatggtcgcgagatctatatctatggcgagcgagtgaaagacgtcactactcatccggcatttcgtaa<br>tgcggctgcgtctgttgcccaactgtacgacgcgctacacaaaccggagatgcaggactctctgtgctggaacac<br>cgacaccggcagcggcggctatacccataaattcttccgcgtggcgaaaagtgccgacgacctgcgccagcaa<br>cgcgatgccatcgctgagtggtcacgcctgagctatggctggatgggccgtaccccagactacaaagctgctttc<br>ggttgtgcactgggcgcgaatGGTCTCAccgggcttttacggtcagttcgagcagaacgcccgtaactggt<br>acacccgtattcaggaaactggcctctactttaaccacgcgattgttaacccaccgatcgatcgtcatttgccgacc<br>gataaagtaaaagacgtttacatcaagctggaaaaagagactgacgccgggattatcgtcagcggtgcgaaagt<br>ggttgccaccaactcggcgctgactcactacaacatgattggcttcggctcggcacaagtaatgggcgaaaaccc<br>ggacttcgcactgatgttcgttgcgccaatggatgccgatggcgtcaaattaatctcccgcgcctcttatgagatgg<br>tcgcgggtgctaccggctcaccgtatgactacccgctctccagccgcttcgatgagaacgatgcgattctggtgat<br>ggataacgtgctgatcccatgggaaaacgtgctgatctaccgcgattttgatcgctgccgtcgctggacgatggaa<br>ggcggtttcgcccgtatgtatccgctgcaagcctgtgtgcgcctggcagtgaaactcgacttcattacggcactgc<br>tgaaaaaatcactcgaatgtaccggcaccctggagttccgtggtgtgcaggccgatctcggtgaagtggtggcgt<br>ggcgcaacaccttctgggcattgagtgactcgatgtgttctgaagcgacgccgtgggtcaacggggcttatttacc<br>ggatcatgccgcactgcaaacctatcgcgtactggcaccaatggcctacgcgaagatcaaaaacattatcgaacg |

| | |
|---|---|
| | caacgttaccagtggcctgatctatctcccttccagtgcccgtgacctgaacaatccgcagatcgaccagtatctg gcgaagtatgtgcgcggttcgaacggtatggatcacgtccagcgcatcaagatcctcaaactgatgtgggatgcc attggcagcgagtttggtggtcgtcacgaactgtatgaaatcaactactctggtagccaggatgagattcgcctgc agtgtctgcgccaggcacaaagctccggcaatatggacaagatgatggcgatggttgatcgctgcctgtcggaat acgaccagaacggctggactgtgccgcacctgcacaacaacgacgatatcaacatgctggataagctgctgaaa taaggatccttcgtcggtcagtttcacctgatttacgtaaaaacccgcttcggcgggtttttgcttttggaggggcag aaagatgaatgactgtcggccattccgggtagcataaccccttggggcctctaaacgggtcttgaggggttttttgt catgtgaggctgccaaaccagatgtcaacacagctacaacgtttatggctagctcagtcctaggtacaatgctagc ggagagccccatagggtggtgtgtaccacccctgatgagtccaaaaggacgaaatggggcctctacaaataattt tgtttaacgtaactacggtacctatgtctcgtttagataaaagtaaagtgattaacagcgcattagagctgcttaatga ggtcggaatcgaaggtttaacaacccgtaaactcgcccagaagctaggtgtagagcagcctacattgtattggcat gtaaaaaataagcgggctttgctcgacgccttagccattgagatgttagataggcaccatactcacttttgccctttta gaaggggaaagctggcaagattttttacgtaataacgctaaaagttttagatgtgctttactaagtcatcgcgatgga gcaaaagtacatttaggtacacggcctacagaaaaacagtatgaaactctcgaaaatcaattagcctttttatgcca acaaggtttttcactagagaatgcattatatgcactcagcgcagtggggcattttactttaggttgcgtattggaagat caagagcatcaagtcgctaaagaagaaagggaaacacctactactgatagtatgccgccattattacgacaagct atcgaattatttgatcaccaaggtgcagagccagccttcttattcggccttgaattgatcatatgcggattagaaaaa caacttaaatgtgaaagtgggtcttaaatcctaactcgaggggaactgccaggcatcaaataaaacgaaaggctc agtcggaagactgggcctttcgttttatctgttgtttgtcggtgaacgctctcctggctggatgcacacactggcttaa gatgacaacgtttatagctagctcagcccttggtacaatgctagcggagagtcataagtctgggctaagcccactg atgagtcgctgaaatgcgacgaaacttatgacctctacaaataattttgtttaatgaaccacgaggccttatgcaatta gatgaacaacgcctgcgctttcgtgacgcaatggccagcctgtcggcagcggtaaatattatcaccaccgaggg cgacgccggacaatgcgggattacggcaacggccgtctgctcggtcacggatacaccaccatcgctgatggtgt gcattaacgccaacagtgcgatgaacccggtttttcagggcaacggtaagttgtgcgtcaacgtcctcaaccatga gcaggaactgatggcacgccacttcgcgggcatgacaggcatggcgatggaagagcgttttagcctctcatgct ggcaaaaaggtccgctggcgcagccggtgctaaaaggttcgctggccagtcttgaaggtgagatccgcgatgtg caggcaattggcacacatctggtgtatctggtggagattaaaaacatcatcctcagtgcagaaggtcacggactta tctactttaaacgccgtttccatccggtgatgctggaaatggaagctgcgatttaaggatcctaactcgaggggaac tgccaggcatcaaataaaacgaaaggctcagtcgaaagactgggcctttcgttttatctgttgtttgtcggtgaacg ctctcctggctgagcagttacagagatgttacgaaccacgcggccgcgattatcaaaaaggatcttcacctagatc cttttaaattaaaaatgaagttttaaatcaatctaaagtatatatgagtaaacttggtctgacagttaccaatgcttaatc agtgaggcacctatctcagcgatctgtctatttcgttcatccatagttgcctgactccccgtcgtgtagataactacga tacgggagggcttaccatctggccccagtgctgcaatgataccgcgggacccacgctcaccggctccagatttat cagcaataaaccagccagccggaagggccgagcgcagaagtggtcctgcaactttatccgcctccatccagtct attaattgttgccgggaagctagagtaagtagttcgccagttaatagtttgcgcaacgttgttgccattgctacaggc atcgtggtgtcacgctcgtcgtttggtatggcttcattcagctccggttcccaacgatcaaggcgagttacatgatcc cccatgttgtgcaaaaaagcggttagctccttcggtcctccgatcgttgtcagaagtaagttggccgcagtgttatc actcatggttatggcagcactgcataattctcttactgtcatgccatccgtaagatgcttttctgtgactggtgagtact caaccaagtcattctgagaatagtgtatgcggcgaccgagttgctcttgcccggcgtcaatacgggataataccgc gccacatagcagaactttaaaagtgctcatcattggaaaacgttcttcggggcgaaaactctcaaggatcttaccg ctgttgagatccagttcgatgtaacccactcgtgcacccaactgatcttcagcatcttttactttcaccagcgtttctgg gtgagcaaaaacaggaaggcaaaatgccgcaaaaaagggaataagggcgacacggaaatgttgaatactcata ctcttcctttttcaatattattgaagcatttatcagggttattgtctcatgagcggatacatatttgaatgtatttagaaaaa taaacaaataggggttccgcgcacatttccccgaaaagtgccacctgggaactatatccggattggcgaatgggt catgaccaaaatcccttaacgtgagttttcgttccactgagcgtcagaccccgtagaaaagatcaaaggatcttc |

| | |
|---|---|
| HpaBC<br>Golden<br>Gate<br>Backbone 2 | ttgagatcctttttttctgcgcgtaatctgctgcttgcaaacaaaaaaaccaccgctaccagcggtggtttgtttgccg<br>gatcaagagctaccaactctttttccgaaggtaactggcttcagcagagcgcagataccaaatactgtccttctagt<br>gtagccgtagttaggccaccacttcaagaactctgtagcaccgcctacatacctcgctctgctaatcctgttaccag<br>tggctgctgccagtggcgataagtcgtgtcttaccgggttggactcaagacgatagttaccggataaggcgcagc<br>ggtcgggctgaacggggggttcgtgcacacagcccagcttggagcgaacgacctacaccgaactgagatacct<br>acagcgtgagctatgagaaagcgccacgcttcccgaagggagaaaggcggacaggtatccggtaagcggca<br>gggtcggaacaggagagcgcacgagggagcttccagggggaaacgcctggtatctttatagtcctgtcgggttt<br>cgccacctctgacttgagcgtcgatttttgtgatgctcgtcaggggggcggagcctatggaaaaacgccagcaac<br>gcggcctttttacggttcctggccttttgctggccttttgctcacatgttctttcctgcgttatcccctgattctgtggata<br>accgtattaccgcctttgagtgagctgataccgctcgccgcagccgaacgactgagcgcagcgagtcagtgagc<br>gaggaagcggaagagcgcctgatgcggtattttctccttacgcatctgtgcggtatttcacaccgcaatggtgcac<br>tctcagtacaatctgctctgatgccgcatagttaagccagtatacactccgctatcgctacgtgactgggtcatggct<br>gcgccccgacacccgccaacacccgctgacgcgccctgacgggcttgtctgctcccggcatccgcttacagac<br>aagctgtgaccatctccgggagctgcatgtgtcagaggttttcaccgtcatcaccgaaacgcgcgaggcagctgc<br>ggtaaagctcatcagcgtggtcgtgaagcgattcacagatgtctgcctgttcatccgcgtccagctcgttgagtttct<br>ccagaagcgttaatgtctggcttctgataaagcgggccatgttaagggcggttttttcctgtttggtcactgatgcctc<br>cgtgtaagggggatttctgttcatgggggtaatgataccgatgaaacgagagaggatgctcacgatacgggttact<br>gatgatgaacgtatatatgagtaaacttggtctgacagaaagcaagctgataaaccgatacaattaaaggctcctttt<br>ggagcctttttttttggagattttcaacatgaaaaaattattattgcggccgcccctgaattcgcatctagactgaactg<br>gccgataattgcagacgaacggcgatacagaccctaatttcacatcatatgacactaatccctatcagtgatagag<br>attgacatccctatcagtgatagagatactgagcacatcagcaggacgcactgaccggagagctgtcaccggat<br>gtgctttccggtctgatgagtccgtgaggacgaaacagcctctacaaataattttgtttaagactttaaagtccaatat<br>gaaaccagaagatttccgcgccagtacccaacgtccgttcaccggggaagagtatctgaaaagcctgcaggatg<br>gtcgcgagatctatatctatggcgagcgagtgaaagacgtcactactcatccggcatttcgtaatgcggctgcgtct<br>gttgcccaactgtacgacgcgctacacaaaccggagatgcaggactctctgtgctggaacaccgacaccggca<br>gcggcggctatacccataaattcttccgcgtggcgaaaagtgccgacgacctgcgccagcaacgcgatgccatc<br>gctgagtggtcacgcctgagctatggctggatgggccgtaccccagactacaaagctgctttcggttgtgcactg<br>ggcgcgaatccggAGAGACCgcttttacggtcagttcgagcagaacgcccgtaactggtacacccgtattc<br>aggaaactggcctctactttaaccacgcgattgttaacccaccgatcgatcgtcatttgccgaccgataaagtaaaa<br>gacgtttacatcaagctggaaaaagagactgacgccgggattatcgtcagcggtgcgaaagtggttgccaccaa<br>ctcggcgctgactcactacaacatgattggcttcggctcggcacaagtaatgggcgaaaacccggacttcgcaG<br>GTCTCActgatgttcgttgcgccaatggatgccgatggcgtcaaattaatctcccgcgcctcttatgagatggt<br>cgcgggtgctaccggctcaccgtatgactacccgctctccagccgcttcgatgagaacgatgcgattctggtgat<br>ggataacgtgctgatcccatgggaaaacgtgctgatctaccgcgattttgatcgctgccgtcgctggacgatggaa<br>ggcggtttcgcccgtatgtatccgctgcaagcctgtgtgcgcctggcagtgaaactcgacttcattacggcactgc<br>tgaaaaaatcactcgaatgtaccggcaccctggagttccgtggtgtgcaggccgatctcggtgaagtggtggcgt<br>ggcgcaacaccttctgggcattgagtgactcgatgtgttctgaagcgacgccgtgggtcaacggggcttatttacc<br>ggatcatgccgcactgcaaacctatcgcgtactggcaccaatggcctacgcgaagatcaaaaacattatcgaacg<br>caacgttaccagtggcctgatctatctcccttccagtgcccgtgacctgaacaatccgcagatcgaccagtatctg<br>gcgaagtatgtgcgcggttcgaacggtatggatcacgtccagcgcatcaagatcctcaaactgatgtgggatgcc<br>attggcagcgagtttggtggtcgtcacgaactgtatgaaatcaactactctggtagccaggatgagattcgcctgc<br>agtgtctgcgccaggcacaaagctccggcaatatggacaagatgatggcgatggttgatcgctgcctgtcggaat<br>acgaccagaacggctggactgtgccgcacctgcacaacaacgacgatatcaacatgctggataagctgctgaaa<br>taaggatccttcgtcggtcagtttcacctgatttacgtaaaaacccgcttcggcgggtttttgcttttggaggggcag<br>aaagatgaatgactgtcggccattccgggtagcataaccccttggggcctctaaacgggtcttgaggggttttttgt<br>catgtgaggctgccaaaccagatgtcaacacagctacaacgtttatggctagctcagtcctaggtacaatgctagc |

| | |
|---|---|
| | ggagagccccatagggtggtgtgtaccacccctgatgagtccaaaaggacgaaatggggcctctacaaataattt<br>tgtttaacgtaactacggtacctatgtctcgtttagataaaagtaaagtgattaacagcgcattagagctgcttaatga<br>ggtcggaatcgaaggtttaacaacccgtaaactcgcccagaagctaggtgtagagcagcctacattgtattggcat<br>gtaaaaaataagcgggctttgctcgacgccttagccattgagatgttagataggcaccatactcacttttgccctta<br>gaaggggaaagctggcaagatttttacgtaataacgctaaaagttttagatgtgctttactaagtcatcgcgatgga<br>gcaaaagtacatttaggtacacggcctacagaaaaacagtatgaaactctcgaaaatcaattagcctttttatgcca<br>acaaggtttttcactagagaatgcattatatgcactcagcgcagtggggcattttactttaggttgcgtattggaagat<br>caagagcatcaagtcgctaaagaagaaagggaaacacctactactgatagtatgccgccattattacgacaagct<br>atcgaattatttgatcaccaaggtgcagagccagccttcttattcggccttgaattgatcatatgcggattagaaaaa<br>caacttaaatgtgaaagtgggtcttaaatcctaactcgaggggaactgccaggcatcaaataaaacgaaaggctc<br>agtcggaagactgggcctttcgttttatctgttgtttgtcggtgaacgctctcctggctggatgcacacactggcttaa<br>gatgacaacgtttatagctagctcagcccttggtacaatgctagcggagagtcataagtctgggctaagcccactg<br>atgagtcgctgaaatgcgacgaaacttatgacctctacaaataattttgtttaatgaaccacgaggccttatgcaatta<br>gatgaacaacgcctgcgctttcgtgacgcaatggccagcctgtcggcagcggtaaatattatcaccaccgaggg<br>cgacgccggacaatgcgggattacggcaacggccgtctgctcggtcacggatacaccaccatcgctgatggtgt<br>gcattaacgccaacagtgcgatgaacccggtttttcagggcaacggtaagttgtgcgtcaacgtcctcaaccatga<br>gcaggaactgatggcacgccacttcgcgggcatgacaggcatggcgatggaagagcgttttagcctctcatgct<br>ggcaaaaaggtccgctggcgcagccggtgctaaaaggttcgctggccagtcttgaaggtgagatccgcgatgtg<br>caggcaattggcacacatctggtgtatctggtggagattaaaaacatcatcctcagtgcagaaggtcacggactta<br>tctactttaaacgccgtttccatccggtgatgctggaaatggaagctgcgatttaaggatcctaactcgaggggaac<br>tgccaggcatcaaataaaacgaaaggctcagtcgaaagactgggcctttcgttttatctgttgtttgtcggtgaacg<br>ctctcctggctgagcagttacagagatgttacgaaccacgcggccgcgattatcaaaaaggatcttcacctagatc<br>cttttaaattaaaaatgaagttttaaatcaatctaaagtatatatgagtaaacttggtctgacagttaccaatgcttaatc<br>agtgaggcacctatctcagcgatctgtctatttcgttcatccatagttgcctgactccccgtcgtgtagataactacga<br>tacgggagggcttaccatctggccccagtgctgcaatgataccgcgggacccacgctcaccggctccagatttat<br>cagcaataaaccagccagccggaagggccgagcgcagaagtggtcctgcaactttatccgcctccatccagtct<br>attaattgttgccgggaagctagagtaagtagttcgccagttaatagtttgcgcaacgttgttgccattgctacaggc<br>atcgtggtgtcacgctcgtcgtttggtatggcttcattcagctccggttcccaacgatcaaggcgagttacatgatcc<br>cccatgttgtgcaaaaaagcggttagctccttcggtcctccgatcgttgtcagaagtaagttggccgcagtgttatc<br>actcatggttatggcagcactgcataattctcttactgtcatgccatccgtaagatgcttttctgtgactggtgagtact<br>caaccaagtcattctgagaatagtgtatgcggcgaccgagttgctcttgcccggcgtcaatacgggataataccgc<br>gccacatagcagaactttaaaagtgctcatcattggaaaacgttcttcggggcgaaaactctcaaggatcttaccg<br>ctgttgagatccagttcgatgtaacccactcgtgcacccaactgatcttcagcatcttttactttcaccagcgtttctgg<br>gtgagcaaaaacaggaaggcaaaatgccgcaaaaaagggaataagggcgacacggaaatgttgaatactcata<br>ctcttcctttttcaatattattgaagcatttatcagggttattgtctcatgagcggatacatatttgaatgtatttagaaaaa<br>taaacaaataggggttccgcgcacatttccccgaaaagtgccacctgggaactatatccggattggcgaatgggt<br>catgaccaaaatcccttaacgtgagttttcgttccactgagcgtcagaccccgtagaaaagatcaaaggatcttc |
| HpaBC Golden Gate Backbone 3 | ttgagatcctttttttctgcgcgtaatctgctgcttgcaaacaaaaaaaccaccgctaccagcggtggtttgtttgccg<br>gatcaagagctaccaactctttttccgaaggtaactggcttcagcagagcgcagataccaaatactgtccttctagt<br>gtagccgtagttaggccaccacttcaagaactctgtagcaccgcctacatacctcgctctgctaatcctgttaccag<br>tggctgctgccagtggcgataagtcgtgtcttaccgggttggactcaagacgatagttaccggataaggcgcagc<br>ggtcgggctgaacggggggttcgtgcacacagcccagcttggagcgaacgacctacaccgaactgagatacct<br>acagcgtgagctatgagaaagcgccacgcttcccgaagggagaaaggcggacaggtatccggtaagcggca<br>gggtcggaacaggagagcgcacgagggagcttccagggggaaacgcctggtatctttatagtcctgtcgggttt<br>cgccacctctgacttgagcgtcgatttttgtgatgctcgtcaggggggcggagcctatggaaaaacgccagcaac<br>gcggcctttttacggttcctggccttttgctggccttttgctcacatgttctttcctgcgttatcccctgattctgtggata |

| | |
|---|---|
| | accgtattaccgcctttgagtgagctgataccgctcgccgcagccgaacgactgagcgcagcgagtcagtgagc<br>gaggaagcggaagagcgcctgatgcggtattttctccttacgcatctgtgcggtatttcacaccgcaatggtgcac<br>tctcagtacaatctgctctgatgccgcatagttaagccagtatacactccgctatcgctacgtgactgggtcatggct<br>gcgccccgacacccgccaacacccgctgacgcgccctgacgggcttgtctgctcccggcatccgcttacagac<br>aagctgtgaccatctccgggagctgcatgtgtcagaggttttcaccgtcatcaccgaaacgcgcgaggcagctgc<br>ggtaaagctcatcagcgtggtcgtgaagcgattcacagatgtctgcctgttcatccgcgtccagctcgttgagtttct<br>ccagaagcgttaatgtctggcttctgataaagcgggccatgttaagggcggttttttcctgtttggtcactgatgcctc<br>cgtgtaagggggatttctgttcatgggggtaatgataccgatgaaacgagagaggatgctcacgatacgggttact<br>gatgatgaacgtatatatgagtaaacttggtctgacagaaagcaagctgataaaccgatacaattaaaggctccttt<br>ggagcctttttttttggagattttcaacatgaaaaaattattattgcggccgcccctgaattcgcatctagactgaactg<br>gccgataattgcagacgaacggcgatacagaccctaatttcacatcatatgacactaatccctatcagtgatagag<br>attgacatccctatcagtgatagagatactgagcacatcagcaggacgcactgaccggagagctgtcaccggat<br>gtgctttccggtctgatgagtccgtgaggacgaaacagcctctacaaataattttgtttaagactttaaagtccaatat<br>gaaaccagaagatttccgcgccagtacccaacgtccgttcaccggggaagagtatctgaaaagcctgcaggatg<br>gtcgcgagatctatatctatggcgagcgagtgaaagacgtcactactcatccggcatttcgtaatgcggctgcgtct<br>gttgcccaactgtatgacgcgctacacaaaccggagatgcaggactctctgtgctggaacaccgacaccggcag<br>cggcggctatacccataaattcttccgcgtggcgaaaagtgccgacgacctgCGCcagcaacgcgatgccat<br>cgctgagtggtcacgcctgagctatggctggatgggccgtaccccagactacaaagctgctttcggttgtgcact<br>gggcgcgaatccgggcttttacggtcagttcgagcagaacgcccgtaactggtacacccgtattcaggaaactg<br>gcctctactttaaccacgcgattgttaacccaccgatcgatcgtcatttgccgaccgataaagtaaaagacgtttaca<br>tcaagctggaaaaagagactgacgccgggattatcgtcagcggtgcgaaagtggttgccaccaactcggcgctg<br>actcactacaacatgattggcttcggggaccggaaggtaatgggcgaaaacccggacttcgcactgaAGAG<br>ACCtgttcgttgcgccaatggatgccgatggcgtcaaattaatctcccgcgcctcttatgagatggtcgcgggtg<br>ctaccggctcaccgtatgactacccgctctccagccgcttcgatgagaacgatgcgattctggtgatggataacgt<br>gctgatcccatgggaaaacgtgctgatctaccgcgattttgatcgctgccgtcgctggacgatggaaggcggtttc<br>gcccgtatgtatccgctgcaagcctgtgtgcgcctggcagtgaaactcgacttcattacggcactgctgaaaaaat<br>cactcgaatgtaccggcaccctggagttccgtggtgtgcaggccgatctcggtgaagtggtggcgtggcgcaac<br>accttctgggcattgagtgactcgatgtgttctgaagcgacgccgtgggtcaacgggGGTCTCAgcttattta<br>ccggatcatgccgcactgcaaacctatcgcgtactggcaccaatggcctacgcgaagatcaaaaacattatcgaa<br>cgcaacgttaccagtggcctgatctatctcccttccagtgcccgtgacctgaacaatccgcagatcgaccagtatct<br>ggcgaagtatgtgcgcggttcgaacggtatggatcacgtccagcgcatcaagatcctcaaactgatgtgggatgc<br>cattggcagcgagtttggtggtcgtcacgaactgtatgaaatcaactactctggtagccaggatgagattcgcctg<br>cagtgtctgcgccaggcacaaagctccggcaatatggacaagatgatggcgatggttgatcgctgcctgtcgga<br>atacgaccagaacggctggactgtgccgcacctgcacaacaacgacgatatcaacatgctggataagctgctga<br>aataaggatccttcgtcggtcagtttcacctgatttacgtaaaaacccgcttcggcgggtttttgcttttggaggggc<br>agaaagatgaatgactgtcggccattccgggtagcataaccccttggggcctctaaacgggtcttgaggggttttt<br>gtcatgtgaggctgccaaaccagatgtcaacacagctacaacgtttatggctagctcagtcctaggtacaatgcta<br>gcggagagccccatagggtggtgtgtaccacccctgatgagtccaaaaggacgaaatggggcctctacaaata<br>attttgtttaacgtaactacggtacctatgtctcgtttagataaaagtaaagtgattaacagcgcattagagctgcttaa<br>tgaggtcggaatcgaaggtttaacaacccgtaaactcgcccagaagctaggtgtagagcagcctacattgtattg<br>gcatgtaaaaaataagcgggctttgctcgacgccttagccattgagatgttagataggcaccatactcacttttgcc<br>ctttagaaggggaaagctggcaagattttttacgtaataacgctaaaagttttagatgtgctttactaagtcatcgcga<br>tggagcaaaagtacatttaggtacacggcctacagaaaaacagtatgaaactctcgaaaatcaattagcctttttat<br>gccaacaaggtttttcactagagaatgcattatatgcactcagcgcagtggggcattttactttaggttgcgtattgga<br>agatcaagagcatcaagtcgctaaagaagaaagggaaacacctactactgatagtatgccgccattattacgaca<br>agctatcgaattatttgatcaccaaggtgcagagccagccttcttattcggccttgaattgatcatatgcggattagaa |

| | |
|---|---|
| | aaacaacttaaatgtgaaagtgggtcttaaatcctaactcgaggggaactgccaggcatcaaataaaacgaaagg<br>ctcagtcggaagactgggcctttcgttttatctgttgtttgtcggtgaacgctctcctggctggatgcacacactggct<br>taagatgacaacgtttatagctagctcagcccttggtacaatgctagcggagagtcataagtctgggctaagccca<br>ctgatgagtcgctgaaatgcgacgaaacttatgacctctacaaataattttgtttaatgaaccacgaggccttatgca<br>attagatgaacaacgcctgcgctttcgtgacgcaatggccagcctgtcggcagcggtaaatattatcaccaccga<br>gggcgacgccggacaatgcgggattacggcaacggccgtctgctcggtcacggatacaccaccatcgctgatg<br>gtgtgcattaacgccaacagtgcgatgaacccggtttttcagggcaacggtaagttgtgcgtcaacgtcctcaacc<br>atgagcaggaactgatggcacgccacttcgcgggcatgacaggcatggcgatggaagagcgttttagcctctca<br>tgctggcaaaaaggtccgctggcgcagccggtgctaaaaggttcgctggccagtcttgaaggtgagatccgcga<br>tgtgcaggcaattggcacacatctggtgtatctggtggagattaaaaacatcatcctcagtgcagaaggtcacgga<br>cttatctactttaaacgccgtttccatccggtgatgctggaaatggaagctgcgatttaaggatcctaactcgaggg<br>gaactgccaggcatcaaataaaacgaaaggctcagtcgaaagactgggcctttcgttttatctgttgtttgtcggtg<br>aacgctctcctggctgagcagttacagagatgttacgaaccacgcggccgcgattatcaaaaaggatcttcaccta<br>gatccttttaaattaaaaatgaagttttaaatcaatctaaagtatatatgagtaaacttggtctgacagttaccaatgctt<br>aatcagtgaggcacctatctcagcgatctgtctatttcgttcatccatagttgcctgactccccgtcgtgtagataact<br>acgatacgggagggcttaccatctggccccagtgctgcaatgataccgcgggacccacgctcaccggctccag<br>atttatcagcaataaaccagccagccggaagggccgagcgcagaagtggtcctgcaactttatccgcctccatcc<br>agtctattaattgttgccgggaagctagagtaagtagttcgccagttaatagtttgcgcaacgttgttgccattgctac<br>aggcatcgtggtgtcacgctcgtcgtttggtatggcttcattcagctccggttcccaacgatcaaggcgagttacat<br>gatcccccatgttgtgcaaaaaagcggttagctccttcggtcctccgatcgttgtcagaagtaagttggccgcagtg<br>ttatcactcatggttatggcagcactgcataattctcttactgtcatgccatccgtaagatgcttttctgtgactggtga<br>gtactcaaccaagtcattctgagaatagtgtatgcggcgaccgagttgctcttgcccggcgtcaatacgggataat<br>accgcgccacatagcagaactttaaaagtgctcatcattggaaaacgttcttcggggcgaaaactctcaaggatct<br>taccgctgttgagatccagttcgatgtaacccactcgtgcacccaactgatcttcagcatcttttactttcaccagcgt<br>ttctgggtgagcaaaaacaggaaggcaaaatgccgcaaaaaagggaataagggcgacacggaaatgttgaata<br>ctcatactcttcctttttcaatattattgaagcatttatcagggttattgtctcatgagcggatacatatttgaatgtatttag<br>aaaaataaacaaataggggttccgcgcacatttccccgaaaagtgccacctgggaactatatccggattggcgaa<br>tgggtcatgaccaaaatcccttaacgtgagttttcgttccactgagcgtcagaccccgtagaaaagatcaaaggat<br>cttc |
| HpaBC Golden Gate Backbone 4 | ttgagatcctttttttctgcgcgtaatctgctgcttgcaaacaaaaaaaccaccgctaccagcggtggtttgtttgccg<br>gatcaagagctaccaactctttttccgaaggtaactggcttcagcagagcgcagataccaaatactgtccttctagt<br>gtagccgtagttaggccaccacttcaagaactctgtagcaccgcctacatacctcgctctgctaatcctgttaccag<br>tggctgctgccagtggcgataagtcgtgtcttaccgggttggactcaagacgatagttaccggataaggcgcagc<br>ggtcgggctgaacggggggttcgtgcacacagcccagcttggagcgaacgacctacaccgaactgagatacct<br>acagcgtgagctatgagaaagcgccacgcttcccgaagggagaaaggcggacaggtatccggtaagcggca<br>gggtcggaacaggagagcgcacgagggagcttccagggggaaacgcctggtatctttatagtcctgtcgggttt<br>cgccacctctgacttgagcgtcgatttttgtgatgctcgtcaggggggcggagcctatggaaaaacgccagcaac<br>gcggcctttttacggttcctggccttttgctggccttttgctcacatgttctttcctgcgttatcccctgattctgtggata<br>accgtattaccgcctttgagtgagctgataccgctcgccgcagccgaacgactgagcgcagcgagtcagtgagc<br>gaggaagcggaagagcgcctgatgcggtattttctccttacgcatctgtgcggtatttcacaccgcaatggtgcac<br>tctcagtacaatctgctctgatgccgcatagttaagccagtatacactccgctatcgctacgtgactgggtcatggct<br>gcgccccgacacccgccaacacccgctgacgcgccctgacgggcttgtctgctcccggcatccgcttacagac<br>aagctgtgaccatctccgggagctgcatgtgtcagaggttttcaccgtcatcaccgaaacgcgcgaggcagctgc<br>ggtaaagctcatcagcgtggtcgtgaagcgattcacagatgtctgcctgttcatccgcgtccagctcgttgagtttct<br>ccagaagcgttaatgtctggcttctgataaagcgggccatgttaagggcggttttttcctgtttggtcactgatgcctc<br>cgtgtaagggggatttctgttcatgggggtaatgataccgatgaaacgagagaggatgctcacgatacgggttact |

| | |
|---|---|
| | gatgatgaacgtatatatgagtaaacttggtctgacagaaagcaagctgataaaccgatacaattaaaggctcctttt<br>ggagcctttttttttggagattttcaacatgaaaaaattattattgcggccgcccctgaattcgcatctagactgaactg<br>gccgataattgcagacgaacggcgatacagaccctaatttcacatcatatgacactaatccctatcagtgatagag<br>attgacatccctatcagtgatagagatactgagcacatcagcaggacgcactgaccggagagctgtcaccggat<br>gtgctttccggtctgatgagtccgtgaggacgaaacagcctctacaaataattttgtttaagactttaaagtccaatat<br>gaaaccagaagatttccgcgccagtacccaacgtccgttcaccggggaagagtatctgaaaagcctgcaggatg<br>gtcgcgagatctatatctatggcgagcgagtgaaagacgtcactactcatccggcatttcgtaatgcggctgcgtct<br>gttgcccaactgtacgacgcgctacacaaaccggagatgcaggactctctgtgctggaacaccgacaccggca<br>gcggcggctatacccataaattcttccgcgtggcgaaaagtgccgacgacctgcgccagcaacgcgatgccatc<br>gctgagtggtcacgcctgagctatggctggatgggccgtaccccagactacaaagctgctttcggttgtgcactg<br>ggcgcgaatccgggcttttacggtcagttcgagcagaacgcccgtaactggtacacccgtattcaggaaactggc<br>ctctactttaaccacgcgattgttaacccaccgatcgatcgtcatttgccgaccgataaagtaaaagacgtttacatc<br>aagctggaaaaagagactgacgccgggattatcgtcagcggtgcgaaagtggttgccaccaactcggcgctga<br>ctcactacaacatgattggcttcggctcggcacaagtaatgggcgaaaacccggacttcgcactgatgttcgttgc<br>gccaatggatgccgatggcgtcaaattaatctcccgcgcctcttatgagatggtcgcgggtgctaccggctcacc<br>gtatgactacccgctctccagccgcttcgatgagaacgatgcgattctggtgatggataacgtgctgatcccatgg<br>gaaaacgtgctgatctaccgcgattttgatcgctgccgtcgctggacgatggaaggcggtttcgcccgtatgtatc<br>cgctgcaagcctgtgtgcgcctggcagtgaaactcgacttcattacggcactgctgaaaaaatcactcgaatgtac<br>cggcaccctggagttccgtggtgtgcaggccgatctcggtgaagtggtggcgtggcgcaacaccttctgggcatt<br>gagtgactcgatgtgttctgaagcgacgccgtgggtcaacggggcttAGAGACCatttaccggatcatgcc<br>gcactgcaaacctatcgcgtactggcaccaatggcctacgcgaagatcaaaaacattatcgaacgcaacgttacc<br>agtggcctgatctatctcccttccagtgcccgtgacctgaacaatccgcagatcgaccagtatctggcgaagtatgt<br>gcgcggttcgaacggtatggatcacgtccagcgcatcaagatcctcaaactgatgtgggatgccattggcagcg<br>agtttggtggtcgtcacgaactgtatgaaatcaactactctggtagccaggatgagattcgcctgcagtgtctgcgc<br>caggcacaaagctccggcaatatggacaagatgatggcgatggttgatcgctgcctgtcggaatacgaccagaa<br>cggctggactgtgccgcacctgcacaacaacgacgatatcaacatgctggataagctgctgaaGGTCTCA<br>ataaggatccttcgtcggtcagtttcacctgatttacgtaaaaacccgcttcggcgggtttttgcttttggaggggca<br>gaaagatgaatgactgtcggccattccgggtagcataaccccttggggcctctaaacgggtcttgaggggttttttg<br>tcatgtgaggctgccaaaccagatgtcaacacagctacaacgtttatggctagctcagtcctaggtacaatgctag<br>cggagagccccatagggtggtgtgtaccacccctgatgagtccaaaaggacgaaatggggcctctacaaataat<br>tttgtttaacgtaactacggtacctatgtctcgtttagataaaagtaaagtgattaacagcgcattagagctgcttaatg<br>aggtcggaatcgaaggtttaacaacccgtaaactcgcccagaagctaggtgtagagcagcctacattgtattggc<br>atgtaaaaaataagcgggctttgctcgacgccttagccattgagatgttagataggcaccatactcacttttgcccttt<br>agaaggggaaagctggcaagatttttacgtaataacgctaaaagttttagatgtgctttactaagtcatcgcgatgg<br>agcaaaagtacatttaggtacacggcctacagaaaaacagtatgaaactctcgaaaatcaattagcctttttatgcc<br>aacaaggtttttcactagagaatgcattatatgcactcagcgcagtggggcattttactttaggttgcgtattggaaga<br>tcaagagcatcaagtcgctaaagaagaaagggaaacacctactactgatagtatgccgccattattacgacaagc<br>tatcgaattatttgatcaccaaggtgcagagccagccttcttattcggccttgaattgatcatatgcggattagaaaaa<br>caacttaaatgtgaaagtgggtcttaaatcctaactcgaggggaactgccaggcatcaaataaaacgaaaggctc<br>agtcggaagactgggcctttcgttttatctgttgtttgtcggtgaacgctctcctggctggatgcacacactggcttaa<br>gatgacaacgtttatagctagctcagcccttggtacaatgctagcggagagtcataagtctgggctaagcccactg<br>atgagtcgctgaaatgcgacgaaacttatgacctctacaaataattttgtttaatgaaccacgaggccttatgcaatta<br>gatgaacaacgcctgcgctttcgtgacgcaatggccagcctgtcggcagcggtaaatattatcaccaccgaggg<br>cgacgccggacaatgcgggattacggcaacggccgtctgctcggtcacggatacaccaccatcgctgatggtgt<br>gcattaacgccaacagtgcgatgaacccggtttttcagggcaacggtaagttgtgcgtcaacgtcctcaaccatga<br>gcaggaactgatggcacgccacttcgcgggcatgacaggcatggcgatggaagagcgttttagcctctcatgct |

| | |
|---|---|
| | ggcaaaaaggtccgctggcgcagccggtgctaaaaggttcgctggccagtcttgaaggtgagatccgcgatgtg<br>caggcaattggcacacatctggtgtatctggtggagattaaaaacatcatcctcagtgcagaaggtcacggactta<br>tctactttaaacgccgtttccatccggtgatgctggaaatggaagctgcgatttaaggatcctaactcgaggggaac<br>tgccaggcatcaaataaaacgaaaggctcagtcgaaagactgggcctttcgttttatctgttgtttgtcggtgaacg<br>ctctcctggctgagcagttacagagatgttacgaaccacgcggccgcgattatcaaaaaggatcttcacctagatc<br>cttttaaattaaaaatgaagttttaaatcaatctaaagtatatatgagtaaacttggtctgacagttaccaatgcttaatc<br>agtgaggcacctatctcagcgatctgtctatttcgttcatccatagttgcctgactccccgtcgtgtagataactacga<br>tacgggagggcttaccatctggccccagtgctgcaatgataccgcgggacccacgctcaccggctccagatttat<br>cagcaataaaccagccagccggaagggccgagcgcagaagtggtcctgcaactttatccgcctccatccagtct<br>attaattgttgccgggaagctagagtaagtagttcgccagttaatagtttgcgcaacgttgttgccattgctacaggc<br>atcgtggtgtcacgctcgtcgtttggtatggcttcattcagctccggttcccaacgatcaaggcgagttacatgatcc<br>cccatgttgtgcaaaaaagcggttagctccttcggtcctccgatcgttgtcagaagtaagttggccgcagtgttatc<br>actcatggttatggcagcactgcataattctcttactgtcatgccatccgtaagatgcttttctgtgactggtgagtact<br>caaccaagtcattctgagaatagtgtatgcggcgaccgagttgctcttgcccggcgtcaatacgggataataccgc<br>gccacatagcagaactttaaaagtgctcatcattggaaaacgttcttcggggcgaaaactctcaaggatcttaccg<br>ctgttgagatccagttcgatgtaacccactcgtgcacccaactgatcttcagcatcttttactttcaccagcgtttctgg<br>gtgagcaaaaacaggaaggcaaaatgccgcaaaaaagggaataagggcgacacggaaatgttgaatactcata<br>ctcttcctttttcaatattattgaagcatttatcagggttattgtctcatgagcggatacatatttgaatgtatttagaaaaa<br>taaacaaataggggttccgcgcacatttccccgaaaagtgccacctgggaactatatccggattggcgaatgggt<br>catgaccaaaatcccttaacgtgagttttcgttccactgagcgtcagaccccgtagaaaagatcaaaggatcttc |
| HpaBC Full Insert Golden Gate Backbone | ttgagatcctttttttctgcgcgtaatctgctgcttgcaaacaaaaaaaccaccgctaccagcggtggtttgtttgccg<br>gatcaagagctaccaactctttttccgaaggtaactggcttcagcagagcgcagataccaaatactgtccttctagt<br>gtagccgtagttaggccaccacttcaagaactctgtagcaccgcctacatacctcgctctgctaatcctgttaccag<br>tggctgctgccagtggcgataagtcgtgtcttaccgggttggactcaagacgatagttaccggataaggcgcagc<br>ggtcgggctgaacggggggttcgtgcacacagcccagcttggagcgaacgacctacaccgaactgagatacct<br>acagcgtgagctatgagaaagcgccacgcttcccgaagggagaaaggcggacaggtatccggtaagcggca<br>gggtcggaacaggagagcgcacgagggagcttccagggggaaacgcctggtatctttatagtcctgtcgggttt<br>cgccacctctgacttgagcgtcgatttttgtgatgctcgtcaggggggcggagcctatggaaaaacgccagcaac<br>gcggcctttttacggttcctggccttttgctggccttttgctcacatgttctttcctgcgttatcccctgattctgtggata<br>accgtattaccgcctttgagtgagctgataccgctcgccgcagccgaacgactgagcgcagcgagtcagtgagc<br>gaggaagcggaagagcgcctgatgcggtattttctccttacgcatctgtgcggtatttcacaccgcaatggtgcac<br>tctcagtacaatctgctctgatgccgcatagttaagccagtatacactccgctatcgctacgtgactgggtcatggct<br>gcgccccgacacccgccaacacccgctgacgcgccctgacgggcttgtctgctcccggcatccgcttacagac<br>aagctgtgaccatctccgggagctgcatgtgtcagaggttttcaccgtcatcaccgaaacgcgcgaggcagctgc<br>ggtaaagctcatcagcgtggtcgtgaagcgattcacagatgtctgcctgttcatccgcgtccagctcgttgagtttct<br>ccagaagcgttaatgtctggcttctgataaagcgggccatgttaagggcggttttttcctgtttggtcactgatgcctc<br>cgtgtaagggggatttctgttcatgggggtaatgataccgatgaaacgagagaggatgctcacgatacgggttact<br>gatgatgaacgtatatatgagtaaacttggtctgacagaaagcaagctgataaaccgatacaattaaaggctcctttt<br>ggagcctttttttttggagattttcaacatgaaaaaattattattgcggccgcccctgaattcgcatctagactgaactg<br>gccgataattgcagacgaacggcgatacagaccctaatttcacatcatatgacactaatccctatcagtgatagag<br>attgacatccctatcagtgatagagatactgagcacatcagcaggacgcactgaccggagagctgtcaccggat<br>gtgctttccggtctgatgagtccgtgaggacgaaacagcctctacaaataattttgtttaagactttaaagtccaatA<br>GAGACCatgaaaccagaagatttccgcgccagtacccaacgtccgttcaccggggaagagtatctgaaaa<br>gcctgcaggatggtcgcgagatctatatctatggcgagcgagtgaaagacgtcactactcatccggcatttcgtaa<br>tgcggctgcgtctgttgcccaactgtacgacgcgctacacaaaccggagatgcaggactctctgtgctggaacac<br>cgacaccggcagcggcggctatacccataaattcttccgcgtggcgaaaagtgccgacgacctgcgccagcaa |

| | |
|---|---|
| | cgcgatgccatcgctgagtggtcacgcctgagctatggctggatgggccgtaccccagactacaaagctgctttc<br>ggttgtgcactgggcgcgaatccgggcttttacggtcagttcgagcagaacgcccgtaactggtacacccgtattc<br>aggaaactggcctctactttaaccacgcgattgttaacccaccgatcgatcgtcatttgccgaccgataaagtaaaa<br>gacgtttacatcaagctggaaaaagagactgacgccgggattatcgtcagcggtgcgaaagtggttgccaccaa<br>ctcggcgctgactcactacaacatgattggcttcggctcggcacaagtaatgggcgaaaacccggacttcgcact<br>gatgttcgttgcgccaatggatgccgatggcgtcaaataatctcccgcgcctcttatgagatggtcgcgggtgcta<br>ccggctcaccgtatgactacccgctctccagccgcttcgatgagaacgatgcgattctggtgatggataacgtgct<br>gatcccatgggaaaacgtgctgatctaccgcgattttgatcgctgccgtcgctggacgatggaaggcggtttcgc<br>ccgtatgtatccgctgcaagcctgtgtgcgcctggcagtgaaactcgacttcattacggcactgctgaaaaaatca<br>ctcgaatgtaccggcaccctggagttccgtggtgtgcaggccgatctcggtgaagtggtggcgtggcgcaacac<br>cttctgggcattgagtgactcgatgtgttctgaagcgacgccgtgggtcaacggggcttatttaccggatcatgcc<br>gcactgcaaacctatcgcgtactggcaccaatggcctacgcgaagatcaaaaacattatcgaacgcaacgttacc<br>agtggcctgatctatctcccttccagtgcccgtgacctgaacaatccgcagatcgaccagtatctggcgaagtatgt<br>gcgcggttcgaacggtatggatcacgtccagcgcatcaagatcctcaaactgatgtgggatgccattggcagcg<br>agtttggtggtcgtcacgaactgtatgaaatcaactactctggtagccaggatgagattcgcctgcagtgtctgcgc<br>caggcacaaagctccggcaatatggacaagatgatggcgatggttgatcgctgcctgtcggaatacgaccagaa<br>cggctggactgtgccgcacctgcacaacaacgacgatatcaacatgctggataagctgctgaaGGTCTCA<br>ataaggatccttcgtcggtcagtttcacctgatttacgtaaaaacccgcttcggcgggtttttgcttttggaggggca<br>gaaagatgaatgactgtcggccattccgggtagcataaccccttggggcctctaaacgggtcttgaggggttttttg<br>tcatgtgaggctgccaaaccagatgtcaacacagctacaacgtttatggctagctcagtcctaggtacaatgctag<br>cggagagccccatagggtggtgtgtaccacccctgatgagtccaaaaggacgaaatggggcctctacaaataat<br>tttgtttaacgtaactacggtacctatgtctcgtttagataaaagtaaagtgattaacagcgcattagagctgcttaatg<br>aggtcggaatcgaaggtttaacaacccgtaaactcgcccagaagctaggtgtagagcagcctacattgtattggc<br>atgtaaaaaataagcgggctttgctcgacgccttagccattgagatgttagataggcaccatactcacttttgcccttt<br>agaaggggaaagctggcaagatttttacgtaataacgctaaaagttttagatgtgctttactaagtcatcgcgatgg<br>agcaaaagtacatttaggtacacggcctacagaaaaacagtatgaaactctcgaaaatcaattagcctttttatgcc<br>aacaaggtttttcactagagaatgcattatatgcactcagcgcagtggggcattttactttaggttgcgtattggaaga<br>tcaagagcatcaagtcgctaaagaagaaagggaaacacctactactgatagtatgccgccattattacgacaagc<br>tatcgaattatttgatcaccaaggtgcagagccagccttcttattcggccttgaattgatcatatgcggattagaaaaa<br>caacttaaatgtgaaagtgggtcttaaatcctaactcgaggggaactgccaggcatcaaataaaacgaaaggctc<br>agtcggaagactgggcctttcgttttatctgttgtttgtcggtgaacgctctcctggctggatgcacacactggcttaa<br>gatgacaacgtttatagctagctcagcccttggtacaatgctagcggagagtcataagtctgggctaagcccactg<br>atgagtcgctgaaatgcgacgaaacttatgacctctacaaataattttgtttaatgaaccacgaggccttatgcaatta<br>gatgaacaacgcctgcgctttcgtgacgcaatggccagcctgtcggcagcggtaaatattatcaccaccgaggg<br>cgacgccggacaatgcgggattacggcaacggccgtctgctcggtcacggatacaccaccatcgctgatggtgt<br>gcattaacgccaacagtgcgatgaacccggtttttcagggcaacggtaagttgtgcgtcaacgtcctcaaccatga<br>gcaggaactgatggcacgccacttcgcgggcatgacaggcatggcgatggaagagcgttttagcctctcatgct<br>ggcaaaaaggtccgctggcgcagccggtgctaaaaggttcgctggccagtcttgaaggtgagatccgcgatgtg<br>caggcaattggcacacatctggtgtatctggtggagattaaaaacatcatcctcagtgcagaaggtcacggactta<br>tctactttaaacgccgtttccatccggtgatgctggaaatggaagctgcgatttaaggatcctaactcgaggggaac<br>tgccaggcatcaaataaaacgaaaggctcagtcgaaagactgggcctttcgttttatctgttgtttgtcggtgaacg<br>ctctcctggctgagcagttacagagatgttacgaaccacgcggccgcgattatcaaaaaggatcttcacctagatc<br>cttttaaattaaaaatgaagttttaaatcaatctaaagtatatatgagtaaacttggtctgacagttaccaatgcttaatc<br>agtgaggcacctatctcagcgatctgtctatttcgttcatccatagttgcctgactccccgtcgtgtagataactacga<br>tacgggagggcttaccatctggccccagtgctgcaatgataccgcgggacccacgctcaccggctccagatttat<br>cagcaataaaccagccagccggaagggccgagcgcagaagtggtcctgcaactttatccgcctccatccagtct |

| | |
|---|---|
| | attaattgttgccgggaagctagagtaagtagttcgccagttaatagtttgcgcaacgttgttgccattgctacaggc atcgtggtgtcacgctcgtcgtttggtatggcttcattcagctccggttcccaacgatcaaggcgagttacatgatcc cccatgttgtgcaaaaaagcggttagctccttcggtcctccgatcgttgtcagaagtaagttggccgcagtgttatc actcatggttatggcagcactgcataattctcttactgtcatgccatccgtaagatgcttttctgtgactggtgagtact caaccaagtcattctgagaatagtgtatgcggcgaccgagttgctcttgcccggcgtcaatacgggataataccgc gccacatagcagaactttaaaagtgctcatcattggaaaacgttcttcggggcgaaaactctcaaggatcttaccg ctgttgagatccagttcgatgtaacccactcgtgcacccaactgatcttcagcatcttttactttcaccagcgtttctgg gtgagcaaaaacaggaaggcaaaatgccgcaaaaaagggaataagggcgacacggaaatgttgaatactcata ctcttcctttttcaatattattgaagcatttatcagggttattgtctcatgagcggatacatatttgaatgtatttagaaaaa taaacaaataggggttccgcgcacatttccccgaaaagtgccacctgggaactatatccggattggcgaatgggt catgaccaaaatcccttaacgtgagttttcgttccactgagcgtcagaccccgtagaaaagatcaaaggatcttc |

Table S2: amino acid sequence and codon optimized DNA sequence for the WT HpaBC

| | |
|---|---|
| WT HpaBC amino acid sequence | MKPEDFRASTQRPFTGEEYLKSLQDGREIYIYGERVKDVTTHPAFRNAAA SVAQLYDALHKPEMQDSLCWNTDTGSGGYTHKFFRVAKSADDLRQQRD AIAEWSRLSYGWMGRTPDYKAAFGCALGANPGFYGQFEQNARNWYTRI QETGLYFNHAIVNPPIDRHLPTDKVKDVYIKLEKETDAGIIVSGAKVVATN SALTHYNMIGFGSAQVMGENPDFALMFVAPMDADGVKLISRASYEMVA GATGSPYDYPLSSRFDENDAILVMDNVLIPWENVLIYRDFDRCRRWTMEG GFARMYPLQACVRLAVKLDFITALLKKSLECTGTLEFRGVQADLGEVVA WRNTFWALSDSMCSEATPWVNGAYLPDHAALQTYRVLAPMAYAKIKNII ERNVTSGLIYLPSSARDLNNPQIDQYLAKYVRGSNGMDHVQRIKILKLMW DAIGSEFGGRHELYEINYSGSQDEIRLQCLRQAQSSGNMDKMMAMVDRC LSEYDQNGWTVPHLHNNDDINMLDKLLK |
| WT HpaBC DNA sequence | ATGAAACCAGAAGATTTCCGCGCCAGTACCCAACGTCCGTTCACCGGG GAAGAGTATCTGAAAAGCCTGCAGGATGGTCGCGAGATCTATATCTAT GGCGAGCGAGTGAAAGACGTCACTACTCATCCGGCATTTCGTAATGCG GCTGCGTCTGTTGCCCAACTGTACGACGCGCTACACAAACCGGAGATG CAGGACTCTCTGTGCTGGAACACCGACACCGGCAGCGGCGGCTATACC CATAAATTCTTCCGCGTGGCGAAAAGTGCCGACGACCTGCGCCAGCAA CGCGATGCCATCGCTGAGTGGTCACGCCTGAGCTATGGCTGGATGGGC CGTACCCCAGACTACAAAGCTGCTTTCGGTTGTGCACTGGGCGCGAAT CCGGGCTTTTACGGTCAGTTCGAGCAGAACGCCCGTAACTGGTACACC CGTATTCAGGAAACTGGCCTCTACTTTAACCACGCGATTGTTAACCCAC CGATCGATCGTCATTTGCCGACCGATAAAGTAAAAGACGTTTACATCA AGCTGGAAAAAGAGACTGACGCCGGGATTATCGTCAGCGGTGCGAAA GTGGTTGCCACCAACTCGGCGCTGACTCACTACAACATGATTGGCTTC GGCTCGGCACAAGTAATGGGCGAAAACCCGGACTTCGCACTGATGTTC GTTGCGCCAATGGATGCCGATGGCGTCAAATTAATCTCCCGCGCCTCTT ATGAGATGGTCGCGGGTGCTACCGGCTCACCGTATGACTACCCGCTCT CCAGCCGCTTCGATGAGAACGATGCGATTCTGGTGATGGATAACGTGC TGATCCCATGGGAAAACGTGCTGATCTACCGCGATTTTGATCGCTGCC GTCGCTGGACGATGGAAGGCGGTTTCGCCCGTATGTATCCGCTGCAAG CCTGTGTGCGCCTGGCAGTGAAACTCGACTTCATTACGGCACTGCTGA AAAAATCACTCGAATGTACCGGCACCCTGGAGTTCCGTGGTGTGCAGG CCGATCTCGGTGAAGTGGTGGCGTGGCGCAACACCTTCTGGGCATTGA |

| | |
|---|---|
| | GTGACTCGATGTGTTCTGAAGCGACGCCGTGGGTCAACGGGGCTTATT<br>TACCGGATCATGCCGCACTGCAAACCTATCGCGTACTGGCACCAATGG<br>CCTACGCGAAGATCAAAAACATTATCGAACGCAACGTTACCAGTGGCC<br>TGATCTATCTCCCTTCCAGTGCCCGTGACCTGAACAATCCGCAGATCGA<br>CCAGTATCTGGCGAAGTATGTGCGCGGTTCGAACGGTATGGATCACGT<br>CCAGCGCATCAAGATCCTCAAACTGATGTGGGATGCCATTGGCAGCGA<br>GTTTGGTGGTCGTCACGAACTGTATGAAATCAACTACTCTGGTAGCCA<br>GGATGAGATTCGCCTGCAGTGTCTGCGCCAGGCACAAAGCTCCGGCAA<br>TATGGACAAGATGATGGCGATGGTTGATCGCTGCCTGTCGGAATACGA<br>CCAGAACGGCTGGACTGTGCCGCACCTGCACAACAACGACGATATCAA<br>CATGCTGGATAAGCTGCTGAAA |

Table S3: Substitutions for the first round of EVOLVEpro predictions

| Double | |
|---|---|
| 1 | A199P_I29V |
| 2 | A199P_H372P |
| 3 | G295T_A199P |
| 4 | L303H_A199P |
| 5 | G295S_A199P |
| Triple | |
| 1 | D354K_A199P_I29V |
| 2 | G295S_A199P_I29V |
| 3 | G295S_A199P_H372P |
| 4 | A199P_H454Q_H372P |
| 5 | L303H_A199P_I29V |
| Quadruple | |
| 1 | G295S_A199P_I29V_H372P |
| 2 | D354K_G295T_A199P_I29V |
| 3 | D354K_A199P_I29V_G122A |
| 4 | V364P_A199P_I29V_T114S |
| 5 | G295T_A199P_H454Q_H372P |
| Quintuple | |
| 1 | R287N_G295T_A199P_G150V_G122A |
| 2 | G295S_A199P_G122A_H454Q_H372P |
| 3 | D354K_A199P_I29V_H454Q_H372P |
| 4 | G295T_A199P_G122A_H454Q_H372P |
| 5 | V364P_A199P_I29V_H454Q_H372P |

Table S4: Substitutions for the second round of EVOLVEpro and Directed Evolution

| Rank | Substitutions |
|---|---|
| ML only | |
| 1 | D354K_A199P_I29V_T114S_H454Q_H372P |

| | | |
|---|---|---|
| 2 | D354K_L303H_A199P_I29V_H454Q_H372P | same as Expanded training data prediction 3 |
| 3 | D354K_P302G_L303H_A199P_H454Q_H372P | |
| 4 | D354K_A199P_I29V_G122A_H454Q_H372P | |
| 5 | D354K_P302G_A199P_I29V_H454Q_H372P | |
| 6 | D354K_A199P_I29V_H454Q_H372P_N481V | |
| Expanded Training Data | | |
| 1 | D354K_S237C_A199P_I29V_H454Q_H372P | |
| 2 | D354K_A199P_H454Q_I438L_H372P | |
| 3 | D354K_L303H_A199P_I29V_H454Q_H372P | same as ML only prediction 2 |
| 4 | D354K_S237C_L303H_A199P_I29V_H372P | |
| 5 | D354K_L303H_A199P_H454Q_I438L_H372P | |
| 6 | D354K_S237C_A199P_I29V_H454Q_I438L | |
| Hybrid Variants | | |
| 1 | S237C_S210L_A211R_Q212L_I157L_N159H_T398A | |
| 2 | T114S_S210L_A211R_Q212L_I157L_N159H | |
| 3 | S237C_S210L_A211R_Q212L_I157L_N159H_I201A | |
| 4 | L303H_S210L_A211R_Q212L_I157L_N159H_I201A | |
| 5 | L303H_S210L_A211R_Q212L_I157L_N159H_T398A | |
| 6 | L303H_T114S_S210L_A211R_Q212L_I157L_N159H_I201A | |
| 7 | L303H_T114S_S210L_A211R_Q212L_I157L_N159H_T398A | |
| 8 | D354K_L303H_T114S_S210L_A211R_Q212L_I157L_N159H_I201A | |
| 9 | D354K_T114S_S210L_A211R_Q212L_I157L_N159H_T398A | |
| 10 | I438L_S210L_A211R_Q212L_I157L_N159H | |
| Directed Evolution | | |
| | R0094H_S0210D_A0211R_Q0212K_A0156S_I0157L_T0398A | |
| | S0210L_A0211R_Q0212L_I0157L_N0159H_T0398A | |
| | S0210L_A0211R_Q0212L_I0157L_N0159H_I0201A | |
| | S0210L_A0211R_Q0212L_I0157L_N0159H | |
